# Transdiagnostic dimensions of psychopathology explain individuals’ unique deviations from normative neurodevelopment in brain structure

**DOI:** 10.1101/2020.06.11.147009

**Authors:** Linden Parkes, Tyler M. Moore, Monica E. Calkins, Philip A. Cook, Matthew Cieslak, David R. Roalf, Daniel H. Wolf, Ruben C. Gur, Raquel E. Gur, Theodore D. Satterthwaite, Danielle S. Bassett

**Author notes:** To whom correspondence should be addressed, Corresponding author: Danielle S. Bassett, Suite 240 Skirkanich Hall, 210 Sth 33rd St, Philadelphia, PA 19104-6321, USA. Authors contributed equally (Theodore D. Satterthwaite, Danielle S. Bassett).

## Abstract

Psychopathology is rooted in neurodevelopment. However, clinical and biological heterogeneity, together with a focus on case-control approaches, have made it difficult to link dimensions of psychopathology to abnormalities of neurodevelopment. Here, using the Philadelphia Neurodevelopmental Cohort, we built normative models of cortical volume and tested whether deviations from these models better predicted psychiatric symptoms compared to raw cortical volume. Specifically, drawing on the *p-factor* hypothesis, we distilled 117 clinical symptom measures into six orthogonal psychopathology dimensions: overall psychopathology, anxious-misery, externalizing disorders, fear, positive psychotic symptoms, and negative psychotic symptoms. We found that multivariate patterns of deviations yielded improved out-of-sample prediction of psychopathology dimensions compared to multivariate patterns of raw cortical volume. We also found that correlations between overall psychopathology and deviations in ventromedial prefrontal, inferior temporal, dorsal anterior cingulate, and insular cortices were stronger than those observed for specific dimensions of psychopathology (e.g., anxious-misery). Notably, these same regions are consistently implicated in a range of putatively distinct disorders. Finally, we performed conventional case-control comparisons of deviations in a group of individuals with depression and a group with attention-deficit hyperactivity disorder (ADHD). We observed spatially overlapping effects between these groups that diminished when controlling for overall psychopathology. Together, our results suggest that modeling cortical brain features as deviations from normative neurodevelopment improves prediction of psychiatric symptoms in out-of-sample testing, and that *p-factor* models of psychopathology may assist in separating biomarkers that are disorder-general from those that are disorder-specific.

## INTRODUCTION

Throughout childhood, adolescence, and young adulthood, the brain undergoes major structural changes that facilitate the emergence of complex behavior and cognition (1,2). Mental disorders often surface during this period (3) and are increasingly understood as resulting from disruptions to normative brain maturation (4,5). While maturational changes are stereotyped at the population level, substantial individual variation also exists (2). The extent to which this individual variation in neurodevelopment may explain psychopathology remains unclear.

Linking abnormalities in neurodevelopment to psychopathology has been limited by several challenges. First, diagnostic nosologies assign individuals with distinct symptom profiles to the same clinical diagnosis, yielding disorder groups with highly heterogenous clinical presentation (6). Second, comorbidity among disorders is high (7–9), rendering it difficult to detect the neural correlates of specific disorders. Third, much of the extant literature has adopted case-control designs that reveal only abnormalities associated with the ‘average’ patient, ignoring the dimensional nature of psychopathology (10). Research linking individuals’ neurodevelopmental alterations with distinct dimensions of psychopathology relevant to multiple disorders is a critical step toward developing diagnostic biomarkers for mental health (11–15).

A promising approach entails examining dimensions of symptoms that cut across diagnostic categories (16). The *p-factor* hypothesis (10,17–19) posits that psychopathology symptoms cluster into latent dimensions including a general factor (known as *p* or ‘overall psychopathology’), which underpins individuals’ tendency to develop all forms of psychopathology, alongside multiple dimensions that describe specific types of psychopathology. This dimensional scoring can be accomplished with a bifactor model (20), which yields specific factors (e.g., externalizing, psychosis) that are orthogonal to the general factor and to each other. Previous research has revealed that such psychopathology dimensions relate to differences in brain structure (19,21–24). However, it remains unclear whether these psychopathology dimensions help to elucidate abnormal neurodevelopment; furthermore, it remains unknown to what extent they can help to dissociate disorder-general from disorder-specific abnormalities.

Here, we evaluated whether dimensions of psychopathology relate to individual differences in deviations from normative neurodevelopment. Using a bifactor model, we modeled overall psychopathology and five specific factors, corresponding to mood and anxiety symptoms, externalizing behavior, fear, positive psychosis symptoms, and negative psychosis symptoms (21,25–27). We integrated these psychopathology dimensions with T1-weighted neuroimaging data using a contemporary machine learning technique known as *normative modeling* (28). Here, a normative model is a statistical model that finds the relationship between age and any brain feature, as well as the variation in this relationship expected in a group of healthy individuals. Then, the brains of individuals who experience psychopathology can be understood with respect to this normative model, allowing identification of regional deviations from normative neurodevelopment for each individual (29–31). This approach allowed us to test whether individuals’ patterns of deviations from normative neurodevelopment were able to predict dimensions of psychopathology in out-of-sample testing. To examine the relative advantages of the normative model, we compared the predictive performance of deviations against predictive performance of raw brain features.

The above framework is applicable to any brain feature that changes reliably with age. Here, we focused on cortical gray matter, indexed via *cortical volume,* which is known to undergo plastic maturation in youth. Cortical volume shows a robust global decrease from childhood to adulthood, reflecting cortical myelination and potentially synaptic pruning (32–34). Prior work has shown widespread non-uniform reductions in cortical volume — as well as thickness and surface area, which together comprise volume — across major depressive disorder (35,36), schizophrenia (37), bipolar disorder (38), and anxiety disorders (39). Across these disorders, overlapping reductions were particularly found in ventromedial prefrontal / medial orbitofrontal cortex (vmPFC/mOFC), inferior temporal, dorsal anterior cingulate (daCC), and insular cortices (39,40).

Here, we sought to understand the link between dimensions of psychopathology from the *p-factor* model and deviations of cortical volume from patterns expected in normative neurodevelopment. We addressed three related questions. First, given that psychopathology may have neurodevelopmental origins, we tested the primary hypothesis that modeling cortical volume according to deviations from normative neurodevelopment would provide improved out-of-sample prediction of psychopathology compared to using raw volume. Second, at the regional level, we expected that overall psychopathology would explain the common abnormalities observed in the case-control literature (35–40); we specifically predicted that greater overall psychopathology would be associated with greater negative deviations (i.e., lower than normative cortical volume) in the vmPFC/mOFC, inferior temporal, daCC, and insular cortices. These regions were selected based on their consistent implication across multiple distinct clinical disorders in both high-powered case-control studies (35–40) and meta-analyses (39). Additionally, supporting the notion that effects in these regions predominantly represent disorder-general rather than disorder-specific biomarkers, we also predicted that correlations between deviations in these regions and overall psychopathology would be stronger than correlations between deviations and the specific dimensions in our model (e.g., externalizing, psychosis). Third, we assessed the extent to which overall psychopathology explained the overlap between whole-brain group-level differences observed for traditional case-control analyses. Specifically, we examined group-level deviations from the normative model in two samples, one with depression and another with attention-deficit hyperactivity disorder (ADHD). We expected that both groups would show spatially correlated patterns of average deviations from normative neurodevelopment. Critically, we hypothesized that the correlation between patterns of average deviations would diminish when overall psychopathology was controlled for in our sample, indicating a lack of sensitivity of a case-control approach to detect disorder-specific biomarkers.

## MATERIALS AND METHODS

### Participants

The institutional review boards of both the University of Pennsylvania and the Children’s Hospital of Philadelphia approved all study procedures. Participants included 1,601 individuals from the Philadelphia Neurodevelopmental Cohort (41), a large community-based study of brain development in youths aged 8 to 22 years. We studied a subset of 1,376 participants, including individuals who were medically healthy and passed stringent quality control benchmarks for the neuroimaging data (see Supplementary Methods).

### Psychopathology dimensions

Details of the dimensional psychopathology model have been published elsewhere (25,26,42) (see Supplementary Methods). Briefly, we used confirmatory bifactor analysis to quantify six orthogonal dimensions of psychopathology, including an overall psychopathology factor common to all symptoms measured herein, and five specific factors: anxious-misery, psychosis-positive, psychosis-negative, externalizing behaviors, and fear (see Table S2 for factor loadings). To ensure normality, psychopathology dimensions were normalized using an inverse normal transformation (43,44).

### Normative modeling

For details on image acquisition, processing, quality control, and derivation of cortical volume see Supplementary Methods. Briefly, regional cortical volume was extracted for each of 400 cortical regions defined by the Schaefer atlas (45). Next, we built normative models to predict regional volume. In order to estimate *normative* neurodevelopmental trajectories, we first split our sample of 1,376 participants into two groups according to the presence or absence of psychiatric history. A total of 408 individuals reported no clinically significant symptoms on any disorder examined, while 983 individuals reported experiencing psychopathology. Next, in order to ensure that our analyses of deviations encompassed the full spectrum of psychopathology — including normal, subclinical, and clinical variance — we randomly sampled 100 of the 408 healthy individuals and combined them with the aforementioned 983 individuals to create a *testing* subset totaling 1,068 individuals. The remaining 308 healthy individuals were designated as our normative *training* subset. Next, for each brain region (*j*), we used Gaussian Process Regression (GPR) to predict cortical volume values from age and sex using the training subset (see Marquand *et al.* (28) for details). A key advantage of this approach is that in addition to fitting potentially non-linear predictions of a brain feature, it also provides regional estimates of the expected variation in the relationship between age and brain features (normative variance) as well as estimates of uncertainty in this variance. Both normative variance and uncertainty are learned by the GPR from the training subset (28). Then, for each participant (*i*) in the testing subset, we generated the predicted cortical volume 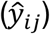 and combined it with the true cortical volume (*y*_*ij*_), the predictive uncertainty (σ_*ij*_), and the normative variance (σ_*nj*_) to create a *z*-score that quantified deviation from normative neurodevelopment (28):

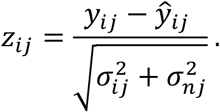

This normative model stands in contrast to alternative approaches, such as brain age models (46), that typically use linear estimates of deviations in age rather than brain features and that do not account for normative variance and estimated uncertainty of deviations (15). In the testing subset, mass univariate application of the normative model yielded a 1,068 × 400 *z*-score deviation matrix, **Z**. Next, we used 10-fold cross-validation to also generate deviations in the training subset. The 308 individuals in the training subset were split into 10 folds, wherein 90% of the subset were used to re-train the GPR in order to generate z-score deviations in the remaining 10%. This process yielded a 308 × 400 *z*-score deviation matrix, **Z**_*cv*_.

### Machine learning prediction models

First, we tested our primary hypothesis that scoring cortical volume as deviations from normative neurodevelopment would yield improved predictive performance compared to using raw volume values. Specifically, columns of **Z**, were taken as multivariate input features to a ridge regression (RR, α = 1) to iteratively predict psychopathology dimensions (*y*). We performed 100 repeats (47) of 10-fold cross-validation scored by root mean squared error (RMSE), mean absolute error (MAE), and the correlation between true and predicted *y* (corr_y). Scoring metrics were averaged over folds. Note, to standardize the interpretation of all scoring metrics as higher scores represent better performance, we flip the sign for RMSE and MAE and examine negative RMSE and negative MAE. Within each fold, we applied principal component analysis (PCA) to reduce the dimensionality of **Z**, to the 9 PCs that explained ≥1% variance in the data (see Figures S5, S6, and S7). Additionally, within each fold, age and sex were controlled for by regressing their effect out of *y* before predicting *y*. Nuisance regression was fit on the training data and applied to the test data to prevent leakage. To assess the significance of prediction performance, for each scoring metric, we averaged over the 100 repeats and compared the corresponding point estimate against a null distribution generated from a model trained on 100,000 random permutations of *y*. The *p*-values were assigned as the proportion of permuted scoring metrics that were greater than or equal to our true point estimates. Then, *p*-values were corrected for multiple comparisons over psychopathology dimensions via the Benjamini and Hochberg False Discovery Rate (FDR, q = 0.05) procedure (48).

The above pipeline generated robust estimates of out-of-sample prediction performance that included significance testing. In order to directly compare predictive performance of normative deviations against raw cortical volume, we repeated this pipeline using a matrix of raw cortical volume values as input features. Note, analysis of raw volume was restricted to the testing subset to maintain equivalent statistical power, and the same number of PCs was used to maintain consistency in the number of input features (see Figures S5, S6, and S7). Together, for each psychopathology dimension and scoring metric, this process yielded two distributions of 100 performance values: one where prediction performance was measured using deviations from a normative model, estimated in an entirely independent sample (*training* subset, n=308), and the other where performance was measured using raw cortical volume as predictors. Critically, as both models included identical nuisance regression applied to *y*, the only difference between models was in the way the cortical features were measured. For each psychopathology dimension and scoring metric, we compared prediction performance across input feature types using an exact test of differences (49).

### Correlations between psychopathology dimensions and regional deviations from normative neurodevelopment

Our machine learning prediction model mapped the relationships between dominant sources of variance in cortical volume (deviations and raw) and psychopathology dimensions. Next, we examined our second hypothesis pertaining to the relative effect sizes of correlations between deviations and overall psychopathology and correlations between deviations and specific dimensions. First, we calculated Pearson correlations between each psychopathology dimension and deviations averaged over subsets of Schaefer parcels that comprised the vmPFC/mOFC, inferior temporal, daCC, and insular areas (see Supplementary Methods and Figure 2). Next, for each region (i.e., daCC), we tested for differences in effect sizes (Pearson’s r) between overall psychopathology and each specific psychopathology dimension using bootstrapping. Specifically, in each of 10,000 bootstrapped samples, we subtracted the absolute effect size for each specific dimension from the absolute effect size for overall psychopathology. Then, upon these distributions of effect size differences, we calculated the 99% confidence interval (CI) and considered overall psychopathology to have yielded a significantly stronger effect if the lower CI bound was >0 (this point corresponded to a Bonferroni-corrected threshold of *p* < 0.01). Second, we supplemented this analysis with a whole-brain mass univariate analysis, wherein Pearson correlations were calculated between each psychopathology dimension and each column in **Z**, (i.e., Schaefer parcels). Here, multiple comparisons were corrected across parcels and psychopathology dimensions (2,400 tests) using the Benjamini and Hochberg False Discovery Rate (FDR, *q* = 0.05) procedure (48) (see Supplementary Methods).

### Case-control comparisons of deviations from normative neurodevelopment

Finally, we tested our third hypothesis that group-level effects derived from case-control designs would be confounded by an insensitivity to disorder-specific effects. We selected a subsample of our testing subset with clinically significant depression (*N* = 144, *Mean age* = 17.62±2.28 years, 33% males) and a subsample with clinically significant ADHD (*N* = 188, *Mean age* = 13.62±3.10, 62% males), two disorders with distinct clinical presentations, and performed case-control analyses. In each group, we excluded participants with comorbid depression and ADHD. We estimated group-level deviations from normative neurodevelopment by calculating regional Cohen’s *d* values comparing the deviations from each group with deviations from the 408 healthy individuals in our sample (i.e., n=308 training subset, **Z**,_*cv*_, and n=100 healthy individuals from the testing subset). Then, to assess correspondence between group-level effects for depression and ADHD, we estimated the spatial (Pearson’s) correlation between regional Cohen’s *d* values. Finally, to examine the extent to which regional variation in Cohen’s *d* values was explained by overall psychopathology, we re-estimated the spatial correlation between regional Cohen’s *d* values after controlling for overall psychopathology.

## RESULTS

### Participants

Sample demographics, including counts of individuals who met diagnostic criteria for lifetime presence of a broad array of disorders, are shown in Table 1 (see also Figure S1 for mean symptom dimensions as a function of diagnostic groups). Males in our sample had significantly higher scores on psychosis-positive (*t* = 2.05, *p* < 0.05, FDR-corrected), psychosis-negative (*t* = 4.23, *p* < 0.05, FDR-corrected), and externalizing (*t* = 5.44, *p* < 0.05, FDR-corrected) factors. Females had significantly higher scores on anxious-misery (*t* = 4.57, *p* < 0.05, FDR-corrected) and fear (*t* = 6.25, *p* < 0.05, FDR-corrected) factors. No significant effect of sex was observed for overall psychopathology. Age correlated significantly with higher overall psychopathology (*r* = 0.33, *p* < 0.05, FDR-corrected) and lower externalizing (*r* = −0.09, *p* < 0.05, FDR-corrected) scores. See Supplementary Results for characterization of trajectories derived from the normative models (i.e., age effects on cortical volume as a function of sex), the effect of age and sex on deviations, as well as sensitivity analyses including a proxy of socioeconomic status (years of maternal education) and parcellation resolution.

**Table 1.**
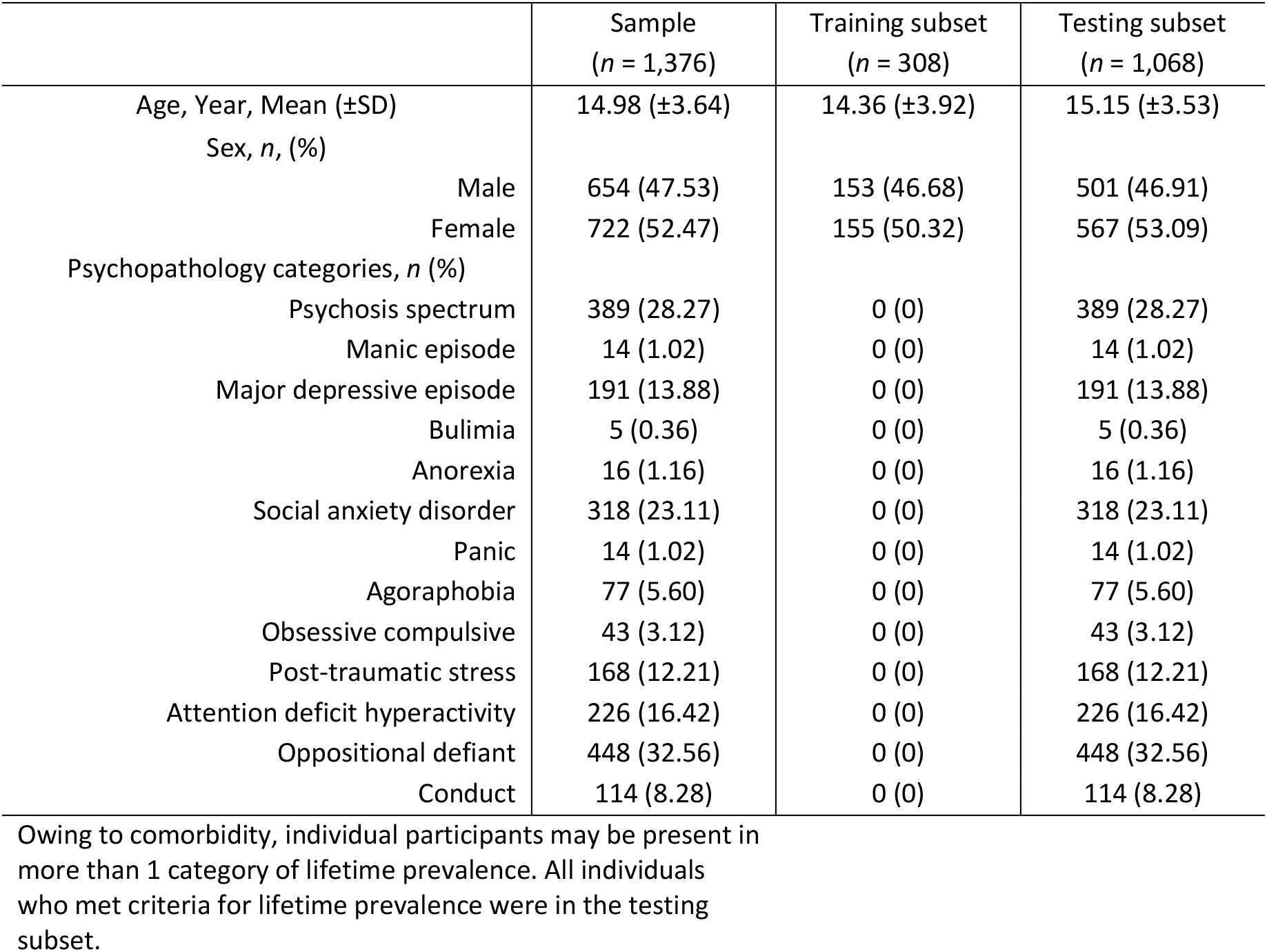
Summary of demographic and psychopathology data

### Modeling deviations from normative neurodevelopment improves prediction of psychopathology dimensions in out-of-sample testing

To address our primary hypothesis that examining deviations from normative models of neurodevelopment would improve predictive performance of dimensions of psychopathology, we compared cross-validated out-of-sample prediction performance when cortical features were scored using deviations to when cortical features were scored using raw volume values. First, our permutation test revealed that we were only able to predict overall psychopathology, psychosis-positive, and fear above chance levels (Figure 1). Critically, for these dimensions, using deviations typically yielded better predictive performance compared to using raw volume across multiple scoring metrics. For example, deviations yielded significantly higher correlations between true and predicted *y* (Figure 1A) and significantly higher negative RMSE (Figure 1B) for overall psychopathology, as well as significantly higher negative MAE for psychosis-positive and fear (Figure 1C). Furthermore, even for dimensions where predictive performance was not above chance levels (i.e., externalizing, psychosis-negative, anxious-misery), we still observed instances where deviations yielded significantly better predictive performance. For example, deviations yielded significantly higher correlations between true and predicted *y* (Figure 1A) for externalizing and significantly higher negative MAE (Figure 1C) for psychosis-negative. In fact, we never observed a psychopathology dimension where (i) raw volume yielded significantly better predictive performance compared to deviations, or (ii) where raw volume yielded above-chance predictive performance, but deviations did not. Thus, in support of hypothesis 1, our results demonstrate that modeling deviations from normative neurodevelopment provided better out-of-sample prediction of psychopathology dimensions compared to using raw cortical volume.

**Figure 1.**
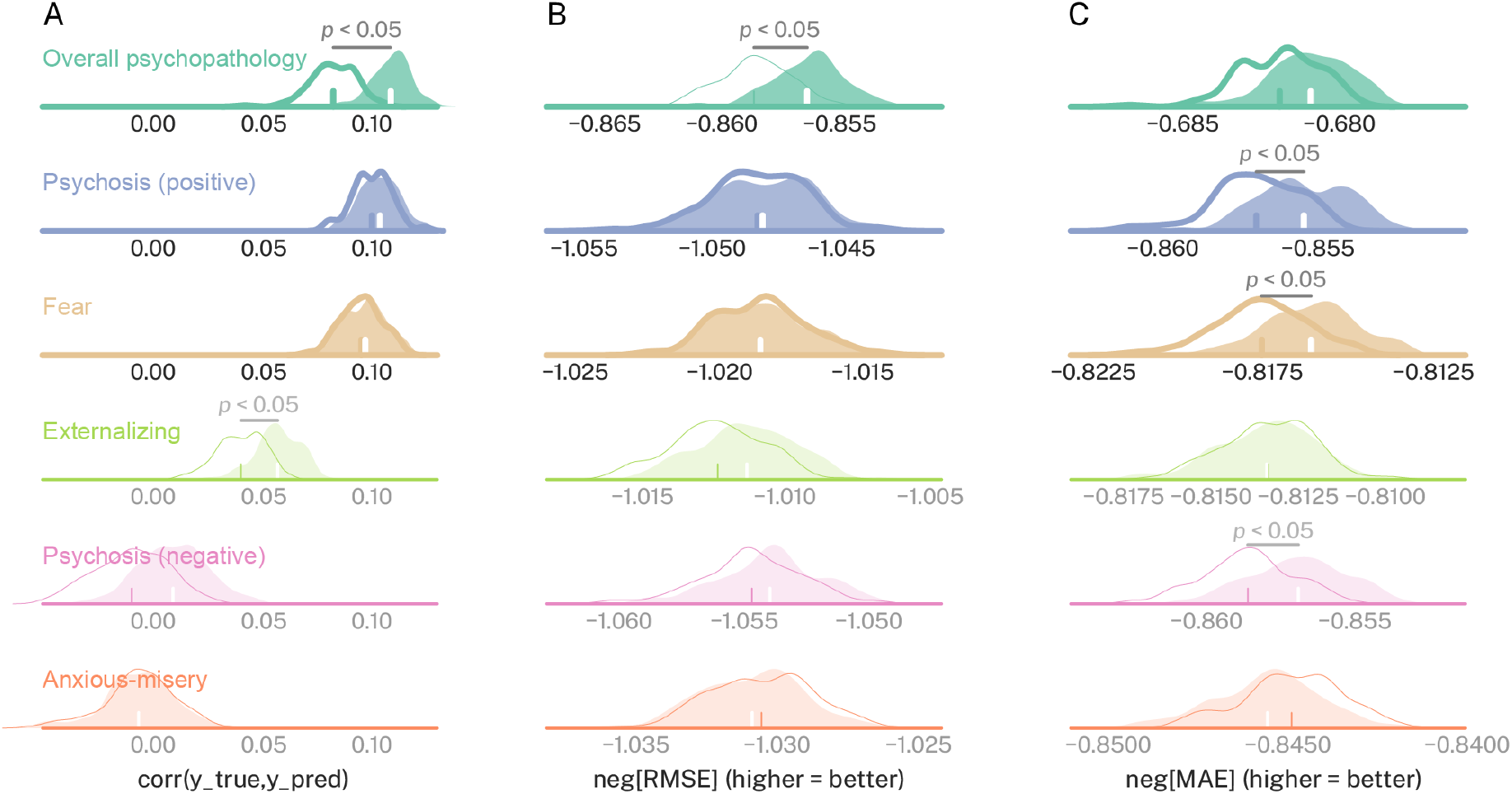
Deviations from normative neurodevelopment yield improved predictive performance of dimensions of psychopathology in out-of-sample testing. Predictive performance for each of six dimensions of psychopathology (rows) as a function of multiple scoring metrics (columns **A-C**). In each subplot two distributions are presented: one that illustrates predictive performance derived from raw cortical volume (white distribution with colored outline), and one that illustrates predictive performance derived from deviations from normative models (colored distribution). Distributions of predictive performance that did not yield above chance performance are shown with partial transparency and lighter stroke. Predictive performance for overall psychopathology, positive psychosis, and fear were above chance levels and are shown with heavier stroke. The scoring metrics based on model error (i.e., RMSE, MAE) are shown with a negative sign so that higher values equal better performance across all scoring metrics. Thus, neg[RMSE] = negative root mean squared error, and neg[MAE] = negative mean absolute error. Differences in scoring metrics between raw cortical volume and deviations were assessed using an exact test of differences. FDR-corrected significant effects are marked with *p* < 0.05.

**Figure 2.**
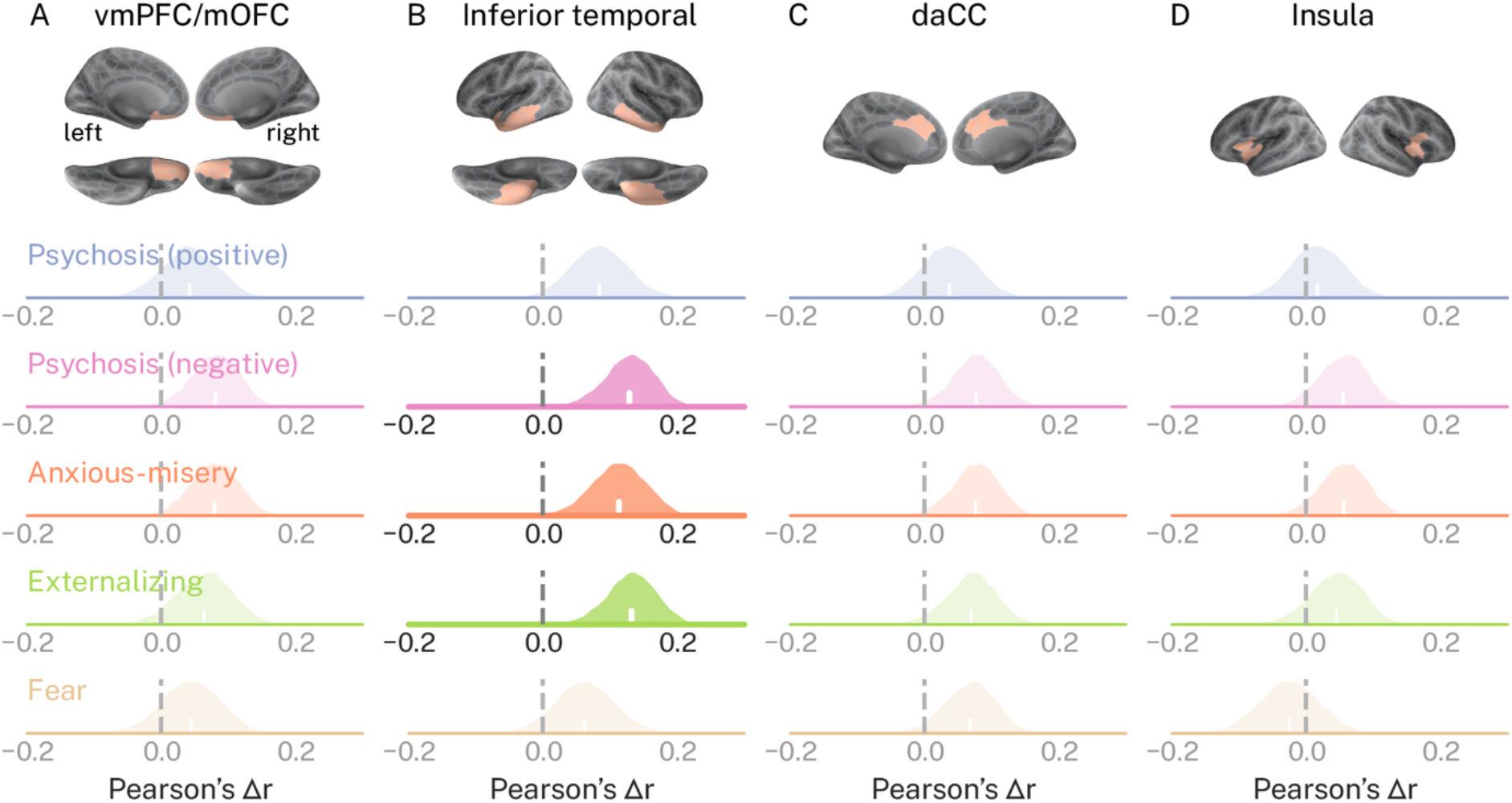
Correlations between overall psychopathology and deviations from normative neurodevelopment are stronger than correlations observed for specific dimensions of psychopathology. In each subplot, distributions of absolute Pearson’s correlation coefficients between each specific psychopathology dimension (rows) and regional deviations (columns) were subtracted from absolute correlations observed for overall psychopathology in the same region. Performing this subtraction 10,000 times across bootstrapped samples generated distributions of effect size differences, Δ*r*. Positive Δ*r* indicates that correlations for overall psychopathology were greater when compared to those observed for the specific dimensions. Δ*r* distributions for which the lower bound of the 99% confidence interval was greater than 0 are shown with heavier stroke and no transparency.

### Psychopathology dimensions explain regional deviations from normative neurodevelopment

Next, we tested our second hypothesis pertaining to the relationships between psychopathology dimensions and deviations from normative neurodevelopment in our *a-priori* regions of interest. First, as expected, we found that overall psychopathology correlated significantly with deviations of the vmPFC/mOFC (*r* = −0.10, *p* < 0.05, FDR-corrected), inferior temporal (*r* = −0.16, *p* < 0.05, FDR-corrected), daCC (*r* = −0.10, *p* < 0.05, FDR-corrected), and insular cortices (*r* = −0.08, *p* < 0.05, FDR-corrected), wherein greater scores corresponded to greater negative deviations. Additionally, the fear dimension significantly correlated with deviations in the inferior temporal (*r* = −0.10, *p* < 0.05, FDR-corrected) and insular cortices (*r* = −0.11, *p* < 0.05, FDR-corrected). Notably, the psychosis-positive, psychosis-negative, anxious-misery, and externalizing dimensions did not yield significant correlations to deviations in any of our regions of interest. Second, we found for each of our regions of interest, correlations between deviations and overall psychopathology were stronger than those observed for specific psychopathology dimensions (Figure 2). However, these effects were only significant for the inferior temporal cortex, where the effect size for overall psychopathology was significantly stronger (at the *p* < 0.01 level) than those observed for psychosis-negative, anxious-misery, and externalizing (Figure 2B). At a more relaxed threshold of *p* < 0.05 (i.e., lower bounds of 95% CI > 0), we found that the effect size for overall psychopathology was significantly larger than those observed for psychosis-positive in the inferior temporal cortex as well as those observed for psychosis-negative and anxious-misery in the mOFC/vmPFC and daCC (Figure S8). No significant effects were observed in the insular cortex (Figure 2 and Figure S8). These results support the notion that abnormalities in regions commonly reported in the case-control literature, particularly the inferior temporal cortex, are strongest for overall psychopathology, suggesting that they may represent disorder-general biomarkers in psychiatry.

Consistent with the above findings, our mass univariate analysis of the 400 Schaefer parcels revealed that greater scores on overall psychopathology was associated with widespread negative deviations across the cortex, including in the vmPFC/mOFC, inferior temporal cortex, daCC, and the insular cortex, albeit to a lesser extent (Figure 3A). Furthermore, compared to overall psychopathology, significant correlations for the specific psychopathology dimensions were sparse, and rarely occurred in our *a-priori* regions of interest (Figure 3B-F). Additionally, consistent with results from our out-of-sample predictive models — that demonstrated we could not predict externalizing, psychosis-negative, or anxious-misery above chance levels — very few significant parcel-wise effects were observed for externalizing (Figure 3D), psychosis-negative (Figure 3E), and the anxious-misery (Figure 3F) dimensions. Finally, our mass univariate analysis revealed a notable consistency in our results, which was that overall psychopathology (Figure 3A), psychosis-positive (Figure 3B), and to a lesser extent fear (Figure 3C), each showed relatively widespread effects in the visual system, and that overall psychopathology showed widespread effects in the somatomotor system.

**Figure 3.**
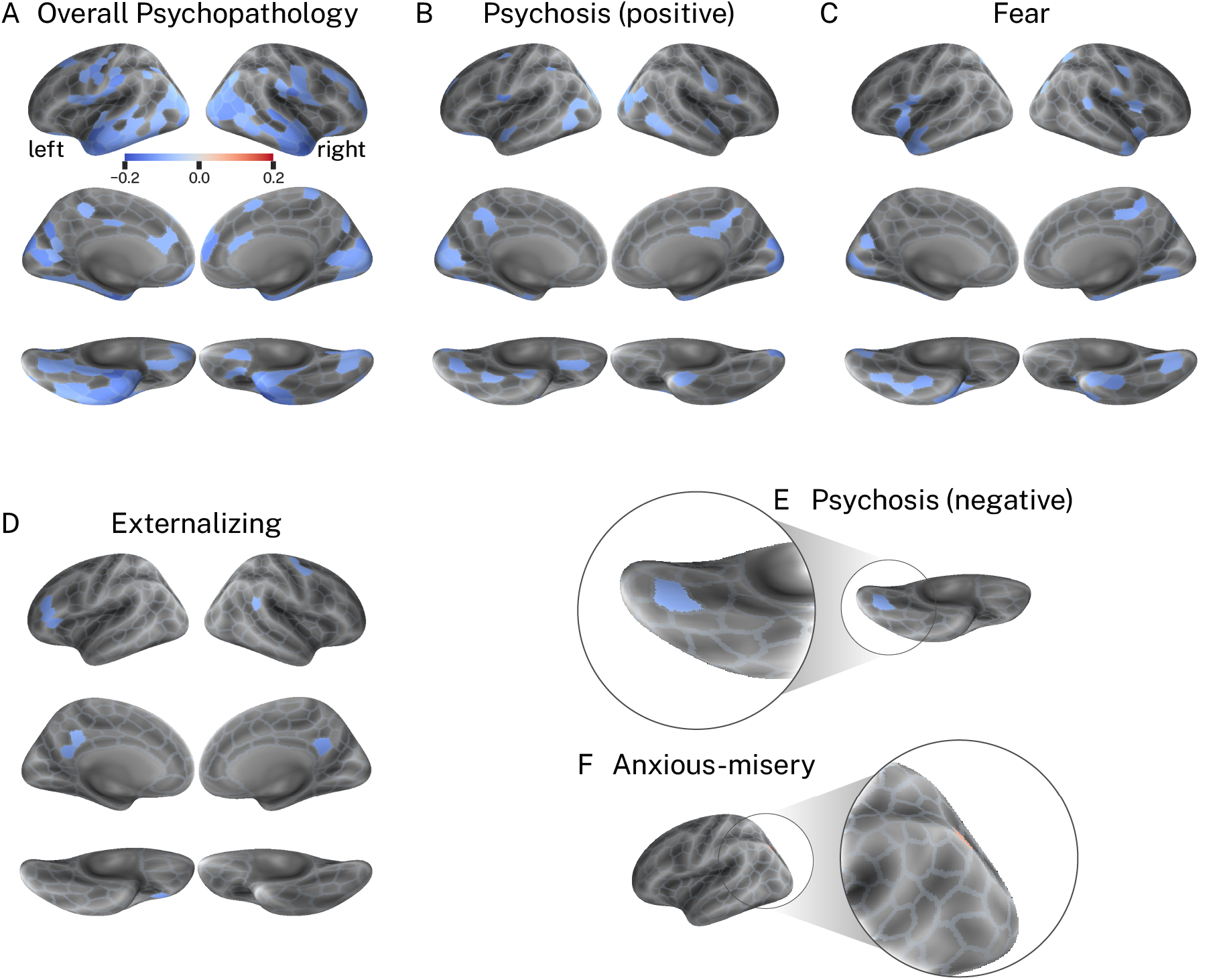
The bivariate relationship between dimensions of psychopathology and deviations from normative neurodevelopment for cortical volume. **A-F**, Significant Pearson’s correlation coefficients between the dimensions of psychopathology and deviations from the normative model. For negative correlations, greater scores on the psychopathology dimension are associated with greater *negative* deviations from normative neurodevelopment. For positive correlations, greater scores on the psychopathology dimension are associated with greater *positive* deviations from normative neurodevelopment (note: this only occurred for anxious-misery; see panel **F**).

### Group effects yield non-specific patterns of deviations from normative neurodevelopment

Above, we demonstrated that overall psychopathology tracked variation in deviations from normative neurodevelopment in regions commonly implicated in the case-control literature. Next, we examined the extent to which overall psychopathology accounted for the spatial overlap between group-level differences observed for case-control analyses conducted in two specific disorders that are conceptualized as dissimilar: depression and ADHD. Figure 4A shows the distribution of Cohen’s *d* values for ADHD and depression for cortical volume with and without controlling for overall psychopathology. For both groups, controlling for overall psychopathology resulted in a significant shift in the Cohen’s *d* distribution towards zero (ADHD, *t* = 31.06, *p* < 0.001; depression, *t* = 31.29, *p* < 0.001). Figure 4B shows the relationship between regional Cohen’s *d* values in each group without controlling for overall psychopathology (*r* = 0.19, *p* < 0.001), while Figure 4C shows the relationship between regional Cohen’s *d* when controlling for the effect of overall psychopathology (*r* = −0.06, *p* = 0.25). Notably, controlling for overall psychopathology reduced the correlation between depression and ADHD to *r* = −0.06; a delta of 0.25. We repeated this analysis using each of the other psychopathology dimensions and found that this relationship was specific to overall psychopathology; when controlling for other dimensions, the spatial correlation delta was, on average, 0.003. Together, these results suggest that the spatial correspondence between group-level deviations for two clinically dissimilar disorders was explained by overall psychopathology.

**Figure 4.**
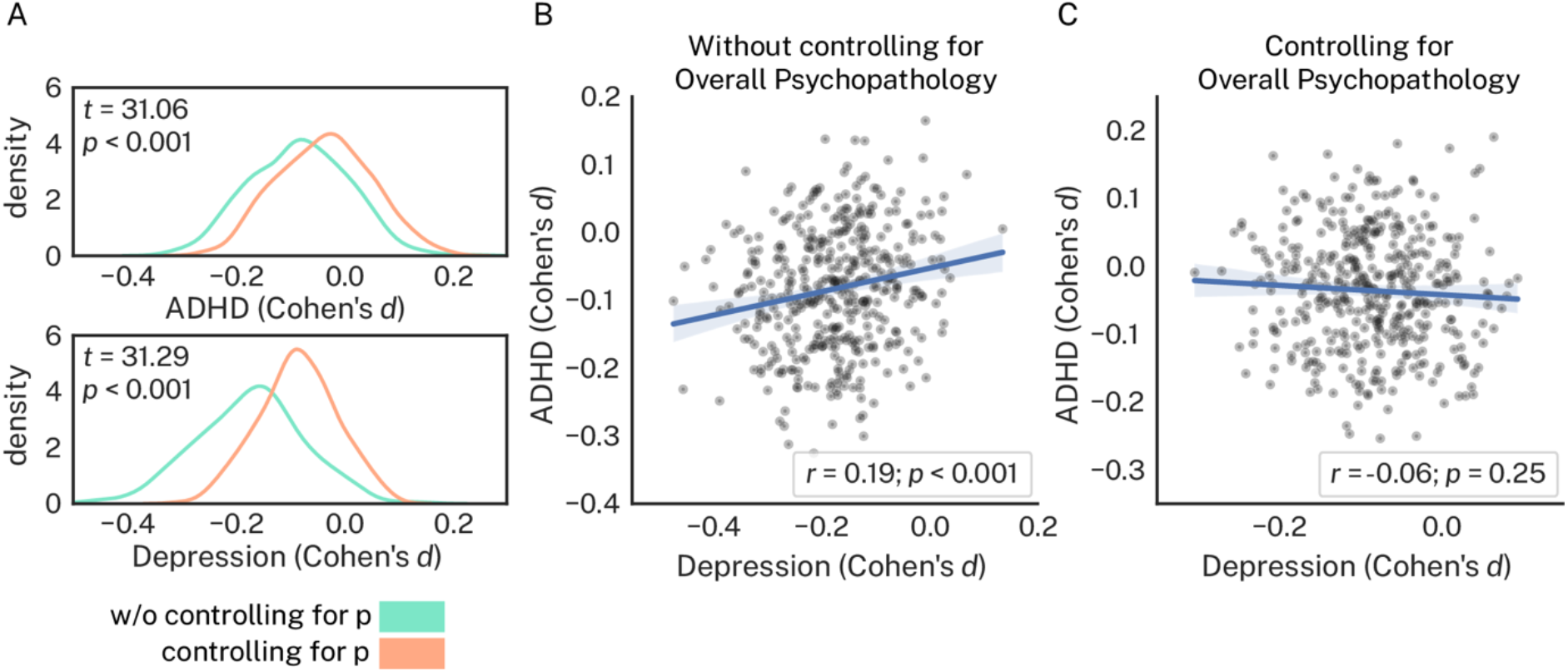
Deviations from normative neurodevelopment in depression and ADHD groups show correlated whole-brain effects confounded by overall psychopathology. Case-control comparisons were conducted examining group differences in deviations between individuals with depression and individuals with ADHD compared to healthy individuals. **A,** Regional Cohen’s *d* values from the ADHD group (top) and depression group (bottom) with and without controlling for overall psychopathology. For both groups, controlling for overall psychopathology resulted in a significant shift in Cohen’s *d* values towards zero. **B**, Regional Cohen’s *d* values from the depression group correlate with regional Cohen’s *d* values from the ADHD group. **C**, Correlations between depression and ADHD groups decrease when controlling for overall psychopathology.

## DISCUSSION

Mental disorders are increasingly viewed as disorders of neurodevelopment (3–5,50). However, heterogeneity in both neurodevelopmental trajectories and symptom profiles have confounded case-control designs and made it difficult to precisely characterize the relationship between abnormalities in neurodevelopment and the symptoms of psychopathology. Here, we showed that modeling cortical brain structure as deviations from normative models of neurodevelopment improved the prediction of dimensions of psychopathology in out-of-sample testing. Furthermore, at the regional level, we demonstrated that overall psychopathology correlated with greater negative deviations in vmPFC/mOFC, inferior temporal, daCC, and insular cortices — all regions previously implicated in case-control literature across a broad spectrum of disorders (35–40) — and that these correlations were stronger than those observed for the specific dimensions of psychopathology in our *p-factor* model. Finally, we found that case-control comparisons between two clinically dissimilar groups (depression and ADHD) and our normative sample showed spatially correlated group differences in deviations that diminished when controlling for overall psychopathology, suggesting that overall psychopathology confounded case-control comparisons. Overall, our results demonstrate that the combination of normative models of neurodevelopment and *p-factor* models of psychopathology not only improve prediction of the symptoms of mental disorder but may also help to tease apart disorder-general from disorder-specific biomarkers in psychiatry.

Previous studies have revealed non-uniform gray matter reductions concentrated in vmPFC/mOFC, inferior temporal, daCC and insular cortices across major depressive, bipolar, schizophrenia, and anxiety disorders (35–40). Here, we found that deviations in each of these regions were implicated most strongly by overall psychopathology. Further, in these regions, significant correlations to deviations were largely absent for our specific psychopathology dimensions. The only exception was fear, for which greater scores corresponded to greater negative deviations in inferior temporal and insular cortices. This effect was broadly recapitulated in our mass univariate analysis, where significant effects were largely absent in the Schaefer parcels that corresponded to our anatomical regions of interest, except in the case of the fear dimension. However, we did observe significant correlations in parcels corresponding to mOFC and inferior temporal cortex for the psychosis-positive dimension, suggesting that positive psychosis symptoms may be associated with abnormally low cortical volume beyond that explained by overall psychopathology. Taken together, our findings suggest that the effects commonly reported in the case-control literature pertaining to the vmPFC/mOFC, inferior temporal, daCC and insular cortices may reflect the general neural correlates of mental disorder more than disorder-specific signatures (39), and that the use of *p-factor* models may assist psychiatry researchers in separating the two.

Beyond our *a-priori* regions of interest, our mass univariate analysis revealed that regions in the visual and somatomotor systems were broadly impacted by overall psychopathology, and that regions in the visual system were also implicated by the psychosis-positive and fear dimensions. This observation is consistent with recent work using functional connectivity data (51,52). Elliot *et al.* (51) showed that overall psychopathology correlated with dysconnectivity between the visual systems and the frontoparietal and default mode systems, and Kebets *et al.* (52) showed that overall psychopathology correlated with dysconnectivity within and between somatomotor and visual systems. Although there are several clear differences between the research of Elliot *et al.* and Kebets *et al.* and the current study, including neuroimaging modality, clinical assessments, statistical methodology, and sample age, the results converge on the idea that disruptions to lower-order brain systems are common across mental disorders. Indeed, given that our sample was, on average, younger than the sample used by Elliot *et al.* (which in turn was younger than that used by Kebets *et al.*), our results suggest that these markers of disorder-general pathophysiology may emerge early during disease progression and persist into adulthood. Datasets covering the lifespan will be critical to testing precise developmental timing effects associated with visual and somatomotor pathophysiology, including the relationship between brain structure and function. Notably, our results, and that of Elliot *et al.* and Kebets *et al.*, suggest that lower-order systems, while important, may be less likely than higher-order systems to yield discriminative utility for mental health.

### Limitations

A limitation of this study is the use of cross-sectional data to model neurodevelopment. It is well documented that individual variability in neurodevelopment occurs at both the inter- and intra-individual level (2), and characterizing the factors that explain the latter will be critical for predicting the emergence of psychopathology over time. Thus, future work should test whether the brain regions identified using our approach explain variance in psychopathology dimensions at follow-up timepoints. Another limitation is the focus on T1-weighted cortical brain features. The underlying white matter pathways change throughout development in parallel with the cortex, giving rise to segregated processing modules that increase the functional efficiency of the brain (53–55). Here, we focused on cortical volume derived from T1-weighted imaging data owing to the robust relationship with age, which is well suited to normative modeling. Future work that builds multimodal normative models of neurodevelopment may provide additional insights into the physiology of psychopathology, which is critical to progressing the field of psychiatry towards personalized medicine.

### Conclusions

Our results represent an important step toward understanding the link between neurodevelopment and psychopathology. We explicitly modeled normative variance in neurodevelopment, allowing us to estimate continuous single-subject neurodevelopmental abnormalities. Combining this approach with a dimensional model of psychopathology allowed us to improve the out-of-sample prediction of psychiatric symptoms compared to raw cortical volume. Further, our work underscores the importance of decoupling specific forms of psychopathology from overall psychopathology. Not doing so may confound the capacity of case-control designs to discover disorder-specific signatures of abnormal neurodevelopment. This confound, in turn, renders case-control designs less likely to yield clinically useful biomarkers in psychiatry. Finally, our work contributes to a growing body of literature demonstrating that, in order to discover neurodevelopmental biomarkers for mental health, psychiatric research could benefit from supplementing examination of the statistical ‘average patient’ with dimensional approaches to psychopathology and brain pathophysiology (14,15,28). Such a neurobiologically-grounded framework may provide a step towards personalized medicine in psychiatry, and ultimately allow for improved outcome for patients.

## ACKNOWLEDGMENTS

The authors acknowledge Dr. Graham Baum for data processing as well as Dr. Urs Braun for valuable discussions during the writing of the manuscript.

## AUTHOR CONTRIBUTIONS

Linden Parkes: Conceptualization, Data curation, Formal analysis, Project administration, Writing - original draft, Writing - review & editing. Tyler M. Moore: Conceptualization, Writing - review & editing. Monica E. Calkins: Conceptualization, Writing - review & editing. Philip A. Cook: Conceptualization, Writing - review & editing. Matthew Cieslak: Conceptualization, Writing - review & editing. David R. Roalf: Conceptualization, Writing - review & editing. Daniel H. Wolf: Conceptualization, Writing - review & editing. Ruben C. Gur: Conceptualization, Writing - review & editing. Raquel E. Gur: Conceptualization, Writing - review & editing. Theodore D. Satterthwaite: Conceptualization, Funding Acquisition, Writing - review & editing. Danielle S. Bassett: Conceptualization, Funding Acquisition, Writing - review & editing.

## CODE AVAILABILITY

All analytic code can be found at: https://github.com/lindenmp/normative_neurodev_cs_t1

## CITATION DIVERSITY STATEMENT

Recent work in several fields of science has identified a bias in citation practices such that papers from women and other minority scholars are under-cited relative to the number of such papers in the field (56–60). Here we sought to proactively consider choosing references that reflect the diversity of the field in thought, form of contribution, gender, race, ethnicity, and other factors. First, we obtained the predicted gender of the first and last author of each reference by using databases that store the probability of a first name being carried by a woman (60,61). By this measure (and excluding self-citations to the first and last authors of our current paper), our references contain 14.55% woman(first)/woman(last), 17.79% man/woman, 24.91% woman/man, and 42.76% man/man. This method is limited in that a) names, pronouns, and social media profiles used to construct the databases may not, in every case, be indicative of gender identity and b) it cannot account for intersex, non-binary, or transgender people. Second, we obtained predicted racial/ethnic category of the first and last author of each reference by databases that store the probability of a first and last name being carried by an author of color (62,63). By this measure (and excluding self-citations), our references contain 11.3% author of color (first)/author of color(last), 24.57% white author/author of color, 17.11% author of color/white author, and 47.02% white author/white author. This method is limited in that a) names and Wikipedia profiles used to make the predictions may not be indicative of racial/ethnic identity, and b) it cannot account for Indigenous and mixed-race authors, or those who may face differential biases due to the ambiguous racialization or ethnicization of their names. We look forward to future work that could help us to better understand how to support equitable practices in science.

## SUPPLEMENTARY METHODS

### SUPPLEMENTARY RESULTS

Table S1. Factor determinacy and Omega-H scores for bifactor model of psychopathology dimensions.

Table S2. Factor loadings from bifactor model of psychopathology dimensions.

**Figure S1. Mean psychopathology symptom dimensions as a function of psychopathology groups.** Groups are the same as those presented in Table 1 in the main text. Only groups where *n*≥50 are shown here.

**Figure S2. Sex effects across psychopathology dimensions in the full sample (n=1,376).**

**Figure S3. Age effects across psychopathology dimensions in the full sample (n=1,376).**

**Figure S4. Age trajectories and model statistics from the normative model. A**, Age trajectories for each sex learned by the normative model. **B**, Out-of-sample standardized mean squared error and explained variance from the normative models run for each region in the Schaefer400 parcellation.

**Figure S5. Variance explained in principal components derived from deviations and raw cortical volume.**

**Figure S6. Coefficients of the principal components derived from deviations from normative neurodevelopment of cortical volume.** The explained variance of each principal component is shown in parentheses.

**Figure S7. Coefficients of the principal components derived from raw cortical volume.** The explained variance of each principal component is shown in parentheses.

**Figure S8. Correlations between overall psychopathology and deviations from normative neurodevelopment are stronger than correlations observed for specific dimensions of psychopathology.**

**Figure S9. Effect of years of maternal education on psychopathology dimensions in the full sample (n=1,376).**

**Figure S10. Prediction model including years of maternal education as a nuisance covariate.**

## SUPPLEMENTARY METHODS

### Participants

From the original 1,601 participants from the Philadelphia Neurodevelopmental Cohort (PNC) (1), 156 were excluded due to the presence of gross radiological abnormalities distorting brain anatomy or due to medical history that might impact brain function; those with a history of psychiatric illness were retained. An additional 17 individuals were excluded due to missing demographic data. Finally, 51 more individuals were excluded because they did not pass rigorous manual and automated quality assurance; one individual was excluded due to corrupted data. This process left a final sample of 1,376 participants. Note that this is a larger sample than studies of normative brain development that have used the PNC; unlike previous reports, we did not exclude based on history of psychiatric illness. Indeed, previous work has illustrated that this broader coverage of the PNC yields prevalence rates of mental disorders consistent with population norms (2).

### Psychopathology phenotypes

In this study, we take a transdiagnostic dimensional approach to assessing variation in the symptoms of mental health (3–6). In particular, we use our recently published extended *p-factor* model (7) based on the GOASSESS interview (8,9). Briefly, the GOASSESS is an abbreviated and modified structured interview derived from the NIMH Genetic Epidemiology Research Branch Kiddie-SADS (10) that covers a wide variety of psychiatric symptomatology such as the occurrence of mood (major depressive episode, mania), anxiety (agoraphobia, generalized anxiety, panic, specific phobia, social phobia, separation anxiety, obsessive compulsive disorder), externalizing behavior (oppositional defiant, attention deficit/hyperactivity, conduct disorder), eating disorder (anorexia, bulimia), and suicidal thoughts and behaviors. GOASSESS was administered by trained and certified assessors. The original model used a combination of exploratory and confirmatory factor analysis to distill the 112 item-level symptoms from the GOASSESS into five orthogonal dimensions of psychopathology. The original model included a factor common to all psychiatric disorders, referred to as overall psychopathology, as well as four specific factors: anxious-misery, psychosis, externalizing behaviors, and fear.

Here, as in our recent work (7), we extended the above *p-factor* model in two ways. First, we included an additional five assessor-rated polytomous items (scored from 0-6, where 0 is ‘absent’ and 6 is ‘severe and psychotic’ or ‘extreme’ from the Scale of Prodromal Symptoms (SOPS) derived from the Structured Interview for Prodromal Syndromes (SIPS (11)) designed to measure the negative/disorganized symptoms of psychosis. These five items were (i) P5 disorganized communication, (ii) N2 avolition, (iii) N3 expression of emotion, (iv) N4 experience of emotions and self, and (v) N6 occupational functioning. Including this additional set of items brought the total number to 117. Second, we split the psychosis factor into two factors, one describing the delusions and hallucinations associated with the psychosis spectrum, which we will refer to as psychosis-positive. The second psychosis factor described disorganized thought, cognitive impairments, and motivational-emotional deficits, which we will refer to as psychosis-negative for simplicity. We used confirmatory factor analysis implemented in Mplus (12) to model five specific factors of psychopathology (anxious-misery, psychosis-positive, psychosis-negative, externalizing behaviors, and fear) as well as one common factor (overall psychopathology). Note that all phenotypes derived from this model are orthogonal to one another.

### Imaging data acquisition

MRI data were acquired on a 3 Tesla Siemens Tim Trio scanner with a 32-channel head coil at the Hospital of the University of Pennsylvania. A 5-min magnetization-prepared, rapid acquisition gradient-echo T1-weighted (MPRAGE) image (TR = 1810ms, TE = 3.51ms, FOV = 180 x 240mm, matrix 256 x 192, voxel resolution of 1mm^3^) was acquired for each participant.

### Imaging data quality control

All T1-weighted images underwent rigorous quality control by highly trained image analysts, see Ref. (13) for details. Briefly, all images were visually inspected and evaluated for the presence of artifacts. Images with gross artifacts were considered unusable; images with some artifacts were flagged as ‘decent’; and images free of artifact were marked as ‘superior’. As mentioned above in the section titled *Participants*, 208 individuals were removed due to unusable imaging data. As a result, 1,139 of our 1,376 participants had T1-weighted images identified as ‘superior’, with the remaining identified as ‘usable’.

### Whole brain parcellation

We summarized cortical volume at the region level. Analyses reported in the main text were conducted using 400 regions covering the cortex that were defined using functional neuroimaging data in a previous study (14). This set of regions is hereafter referred as the Schaefer400 parcellation. However, there exists a plethora of parcellations in neuroimaging research that vary in their construction. In light of this diversity, we sought to confirm that our results were not driven by choice of brain parcellation, and thus we repeated our analyses using a separate parcellation wherein boundaries were defined according to neuroanatomy rather than function (15) and that included 463 regions (Lausanne463). For each parcellation, brain features were generated for each participant as described in the following sections.

### Structural image processing

Structural image processing used tools included in ANTs (16). Structural images were processed in participant’s native space using the following procedure: brain extraction, N4 bias field correction (17), Atropos tissue segmentation (18), and SyN diffeomorphic registration (19,20). Regional estimates of cortical volume were extracted from every participant’s native space data using the following procedure. First, we created a custom adolescent template and tissue priors using data from 140 PNC participants, balanced for age and sex. A custom template minimizes registration bias and maximizes sensitivity to detect regional effects that can be impacted by registration error. Second, the Schaefer400 parcellation was generated in this template space. Third, non-linear registration warps were generated that mapped each participant’s structural scan to template space, and the inverse of these warps were applied to the Schaefer400 parcellation to generate participant-specific parcellations. Participant-specific Lausanne463 parcellations were generated in native space data using code available online: https://github.com/mattcieslak/easy_lausanne. Fourth, each participant’s Schaefer400 parcellation was masked by a cortical gray matter mask from ANTs, also in participant’s native space. Finally, regional cortical volume estimates were extracted using these participant-specific parcellations (Schafer400, Lausanne463) as the count of the number of voxels in each parcel.

### Regional analysis of a priori regions of interest

In the main text, we reported analyzing relationships between psychopathology dimensions and deviations in the vmPFC/mOFC, inferior temporal, daCC and insular cortices. In particular, we referenced the fact that these regions were generated by averaging deviations of subsets of Schaefer parcels (see Figure 2). Here, we provide a full list of the names of these parcels.

The vmPFC/mOFC was comprised of the following parcels:

17Networks_LH_Limbic_OFC_1, 17Networks_LH_Limbic_OFC_2,
17Networks_LH_Limbic_OFC_3, 17Networks_LH_Limbic_OFC_4,
17Networks_LH_Limbic_OFC_5, 17Networks_LH_SalVentAttnB_OFC_1,
17Networks_RH_Limbic_OFC_1, 17Networks_RH_Limbic_OFC_2,
17Networks_RH_Limbic_OFC_3, 17Networks_RH_Limbic_OFC_4,
17Networks_RH_Limbic_OFC_5, 17Networks_RH_Limbic_OFC_6.

The inferior temporal cortex was comprised of the following parcels:

17Networks_LH_Limbic_TempPole_1, 17Networks_LH_Limbic_TempPole_2,
17Networks_LH_Limbic_TempPole_3, 17Networks_LH_Limbic_TempPole_4,
17Networks_LH_Limbic_TempPole_5, 17Networks_LH_Limbic_TempPole_6,
17Networks_LH_Limbic_TempPole_7, 17Networks_LH_ContB_Temp_1,
17Networks_LH_DefaultB_Temp_1, 17Networks_LH_DefaultB_Temp_2,
17Networks_LH_DefaultB_Temp_3, 17Networks_LH_DefaultB_Temp_4,
17Networks_LH_DefaultB_Temp_5, 17Networks_LH_DefaultB_Temp_6,
17Networks_RH_Limbic_TempPole_1, 17Networks_RH_Limbic_TempPole_2,
17Networks_RH_Limbic_TempPole_3, 17Networks_RH_Limbic_TempPole_4,
17Networks_RH_Limbic_TempPole_5, 17Networks_RH_Limbic_TempPole_6,
17Networks_RH_ContB_Temp_1, 17Networks_RH_ContB_Temp_2,
17Networks_RH_DefaultA_Temp_1, 17Networks_RH_DefaultB_Temp_1,
17Networks_RH_DefaultB_Temp_2, 17Networks_RH_DefaultB_AntTemp_1.

The daCC was comprised of the following parcels:

17Networks_LH_SalVentAttnB_PFCmp_1, 17Networks_LH_DefaultA_PFCm_6,
17Networks_LH_ContA_Cinga_1, 17Networks_RH_SalVentAttnB_PFCmp_1,
17Networks_RH_SalVentAttnB_PFCmp_2, 17Networks_RH_DefaultA_PFCm_6,
17Networks_RH_ContA_Cinga_1.

The insular cortex was comprised of the following parcels:

17Networks_LH_SalVentAttnA_Ins_1, 17Networks_LH_SalVentAttnA_Ins_2,
17Networks_LH_SalVentAttnA_Ins_3, 17Networks_LH_SalVentAttnA_Ins_4,
17Networks_LH_SalVentAttnA_Ins_5, 17Networks_LH_SalVentAttnA_Ins_6
17Networks_RH_SalVentAttnA_Ins_1, 17Networks_RH_SalVentAttnA_Ins_2,
17Networks_RH_SalVentAttnA_Ins_3, 17Networks_RH_SalVentAttnA_Ins_4,
17Networks_RH_SalVentAttnA_Ins_5, 17Networks_RH_SalVentAttnA_Ins_6,
17Networks_RH_SalVentAttnA_Ins_7.

## SUPPLEMENTARY RESULTS

### Dimensional measures of psychopathology

Below we illustrate model statistics (Table S1) and factor loadings (Table S2) for our bifactor model of psychopathology.

**Table S1.** Factor determinacy and Omega-H scores for the bifactor model of psychopathology dimensions.

| Item | General<br>(*p'*) | Psychosis-positive | Psychosis-negative | Anxious-misery | Externalizing | Fear |
| --- | --- | --- | --- | --- | --- | --- |
| Factor<br>determinacy | 0.9927 | 0.9600 | 0.9683 | 0.9548 | 0.9661 | 0.9502 |
| Omega-H<sub>subscale</sub> | 0.9213 | 0.0154 | 0.0056 | 0.0004 | 0.0276 | 0.0192 |

**Table S2.** Factor loadings from the bifactor model of psychopathology dimensions.

| Item | Loadings |  |  |  |  |  |
| --- | --- | --- | --- | --- | --- | --- |
|  | General<br>(*p'*) | Psychosis-positive | Psychosis-negative | Anxious-misery | Externalizing | Fear |
| psy001 | **0.657** | **0.442** | 0.000 | 0.000 | 0.000 | 0.000 |
| psy029 | **0.606** | **0.411** | 0.000 | 0.000 | 0.000 | 0.000 |
| psy050 | **0.632** | **0.220** | 0.000 | 0.000 | 0.000 | 0.000 |
| psy060 | **0.666** | **0.316** | 0.000 | 0.000 | 0.000 | 0.000 |
| psy070 | **0.637** | **0.285** | 0.000 | 0.000 | 0.000 | 0.000 |
| psy071 | **0.721** | **0.187** | 0.000 | 0.000 | 0.000 | 0.000 |
| sip003 | **0.598** | **0.522** | 0.000 | 0.000 | 0.000 | 0.000 |
| sip004 | **0.422** | **0.616** | 0.000 | 0.000 | 0.000 | 0.000 |
| sip005 | **0.593** | **0.605** | 0.000 | 0.000 | 0.000 | 0.000 |
| sip006 | **0.557** | **0.559** | 0.000 | 0.000 | 0.000 | 0.000 |
| sip007 | **0.584** | **0.608** | 0.000 | 0.000 | 0.000 | 0.000 |
| sip008 | **0.519** | **0.628** | 0.000 | 0.000 | 0.000 | 0.000 |
| sip009 | **0.615** | **0.502** | 0.000 | 0.000 | 0.000 | 0.000 |
| sip010 | **0.437** | **0.666** | 0.000 | 0.000 | 0.000 | 0.000 |
| sip011 | **0.623** | **0.607** | 0.000 | 0.000 | 0.000 | 0.000 |
| sip012 | **0.639** | **0.596** | 0.000 | 0.000 | 0.000 | 0.000 |
| sip013 | **0.605** | **0.593** | 0.000 | 0.000 | 0.000 | 0.000 |
| sip014 | **0.715** | **0.489** | 0.000 | 0.000 | 0.000 | 0.000 |
| sip027 | **0.487** | 0.000 | **0.288** | 0.000 | 0.000 | 0.000 |
| sip028 | **0.517** | 0.000 | **0.305** | 0.000 | 0.000 | 0.000 |
| sip032 | **0.758** | 0.000 | **0.188** | 0.000 | 0.000 | 0.000 |
| sip033 | **0.681** | 0.000 | **0.205** | 0.000 | 0.000 | 0.000 |
| sip038 | **0.396** | 0.000 | **0.795** | 0.000 | 0.000 | 0.000 |
| sip039 | **0.483** | 0.000 | **0.631** | 0.000 | 0.000 | 0.000 |
| SIP030 | **0.524** | 0.000 | **0.383** | 0.000 | 0.000 | 0.000 |
| SIP035 | **0.714** | 0.000 | **0.302** | 0.000 | 0.000 | 0.000 |
| SIP037 | **0.387** | 0.000 | **0.395** | 0.000 | 0.000 | 0.000 |
| SIP041 | **0.459** | 0.000 | **0.846** | 0.000 | 0.000 | 0.000 |
| SIP043 | **0.496** | 0.000 | **0.678** | 0.000 | 0.000 | 0.000 |
| SIP001 | **0.461** | 0.000 | **0.328** | 0.000 | 0.000 | 0.000 |
| add011 | **0.473** | 0.000 | 0.000 | 0.000 | **0.745** | 0.000 |
| add012 | **0.458** | 0.000 | 0.000 | 0.000 | **0.749** | 0.000 |
| add013 | **0.490** | 0.000 | 0.000 | 0.000 | **0.596** | 0.000 |
| add014 | **0.442** | 0.000 | 0.000 | 0.000 | **0.606** | 0.000 |
| add015 | **0.499** | 0.000 | 0.000 | 0.000 | **0.565** | 0.000 |
| add016 | **0.510** | 0.000 | 0.000 | 0.000 | **0.678** | 0.000 |
| add020 | **0.497** | 0.000 | 0.000 | 0.000 | **0.543** | 0.000 |
| add021 | **0.448** | 0.000 | 0.000 | 0.000 | **0.599** | 0.000 |
| add022 | **0.468** | 0.000 | 0.000 | 0.000 | **0.603** | 0.000 |
| agr001 | **0.611** | 0.000 | 0.000 | 0.000 | 0.000 | **0.474** |
| agr002 | **0.635** | 0.000 | 0.000 | 0.000 | 0.000 | **0.489** |
| agr003 | **0.651** | 0.000 | 0.000 | 0.000 | 0.000 | **0.421** |
| agr004 | **0.550** | 0.000 | 0.000 | 0.000 | 0.000 | **0.422** |
| agr005 | **0.523** | 0.000 | 0.000 | 0.000 | 0.000 | **0.469** |
| agr006 | **0.620** | 0.000 | 0.000 | 0.000 | 0.000 | **0.457** |
| agr007 | **0.621** | 0.000 | 0.000 | 0.000 | 0.000 | **0.286** |
| agr008 | **0.621** | 0.000 | 0.000 | 0.000 | 0.000 | **0.453** |
| cdd001 | **0.573** | 0.000 | 0.000 | 0.000 | **0.407** | 0.000 |
| cdd002 | **0.548** | 0.000 | 0.000 | 0.000 | **0.219** | 0.000 |
| cdd003 | **0.621** | 0.000 | 0.000 | 0.000 | **0.462** | 0.000 |
| cdd004 | **0.468** | 0.000 | 0.000 | 0.000 | **0.334** | 0.000 |
| cdd005 | **0.606** | 0.000 | 0.000 | 0.000 | **0.477** | 0.000 |
| cdd006 | **0.613** | 0.000 | 0.000 | 0.000 | **0.384** | 0.000 |
| cdd007 | **0.635** | 0.000 | 0.000 | 0.000 | **0.372** | 0.000 |
| cdd008 | **0.637** | 0.000 | 0.000 | 0.000 | **0.348** | 0.000 |
| dep001 | **0.760** | 0.000 | 0.000 | **0.220** | 0.000 | 0.000 |
| dep002 | **0.724** | 0.000 | 0.000 | **0.187** | 0.000 | 0.000 |
| dep004 | **0.791** | 0.000 | 0.000 | **0.031** | 0.000 | 0.000 |
| dep006 | **0.775** | 0.000 | 0.000 | **0.034** | 0.000 | 0.000 |
| gad001 | **0.506** | 0.000 | 0.000 | **0.377** | 0.000 | 0.000 |
| gad002 | **0.554** | 0.000 | 0.000 | **0.404** | 0.000 | 0.000 |
| man001 | **0.743** | 0.000 | 0.000 | **-0.517** | 0.000 | 0.000 |
| man002 | **0.744** | 0.000 | 0.000 | **-0.567** | 0.000 | 0.000 |
| man003 | **0.732** | 0.000 | 0.000 | **-0.523** | 0.000 | 0.000 |
| man004 | **0.771** | 0.000 | 0.000 | **-0.456** | 0.000 | 0.000 |
| man005 | **0.767** | 0.000 | 0.000 | **-0.460** | 0.000 | 0.000 |
| man006 | **0.689** | 0.000 | 0.000 | **-0.487** | 0.000 | 0.000 |
| man007 | **0.808** | 0.000 | 0.000 | **-0.241** | 0.000 | 0.000 |
| ocd001 | **0.844** | 0.000 | 0.000 | **0.197** | 0.000 | 0.000 |
| ocd002 | **0.807** | 0.000 | 0.000 | **0.125** | 0.000 | 0.000 |
| ocd003 | **0.709** | 0.000 | 0.000 | **0.209** | 0.000 | 0.000 |
| ocd004 | **0.826** | 0.000 | 0.000 | **0.060** | 0.000 | 0.000 |
| ocd005 | **0.822** | 0.000 | 0.000 | **0.115** | 0.000 | 0.000 |
| ocd006 | **0.843** | 0.000 | 0.000 | **0.107** | 0.000 | 0.000 |
| ocd007 | **0.665** | 0.000 | 0.000 | **0.143** | 0.000 | 0.000 |
| ocd008 | **0.766** | 0.000 | 0.000 | **0.131** | 0.000 | 0.000 |
| ocd011 | **0.712** | 0.000 | 0.000 | **0.196** | 0.000 | 0.000 |
| ocd012 | **0.721** | 0.000 | 0.000 | **0.134** | 0.000 | 0.000 |
| ocd013 | **0.699** | 0.000 | 0.000 | **0.119** | 0.000 | 0.000 |
| ocd014 | **0.763** | 0.000 | 0.000 | **0.061** | 0.000 | 0.000 |
| ocd015 | **0.732** | 0.000 | 0.000 | **0.092** | 0.000 | 0.000 |
| ocd016 | **0.714** | 0.000 | 0.000 | **0.150** | 0.000 | 0.000 |
| ocd017 | **0.719** | 0.000 | 0.000 | **0.090** | 0.000 | 0.000 |
| ocd018 | **0.629** | 0.000 | 0.000 | **0.095** | 0.000 | 0.000 |
| ocd019 | **0.561** | 0.000 | 0.000 | **0.073** | 0.000 | 0.000 |
| odd001 | **0.588** | 0.000 | 0.000 | 0.000 | **0.436** | 0.000 |
| odd002 | **0.573** | 0.000 | 0.000 | 0.000 | **0.515** | 0.000 |
| odd003 | **0.532** | 0.000 | 0.000 | 0.000 | **0.568** | 0.000 |
| odd005 | **0.553** | 0.000 | 0.000 | 0.000 | **0.486** | 0.000 |
| odd006 | **0.634** | 0.000 | 0.000 | 0.000 | **0.397** | 0.000 |
| pan001 | **0.621** | 0.000 | 0.000 | **0.275** | 0.000 | 0.000 |
| pan003 | **0.692** | 0.000 | 0.000 | **0.156** | 0.000 | 0.000 |
| pan004 | **0.779** | 0.000 | 0.000 | **0.159** | 0.000 | 0.000 |
| phb001 | **0.276** | 0.000 | 0.000 | 0.000 | 0.000 | **0.309** |
| phb002 | **0.340** | 0.000 | 0.000 | 0.000 | 0.000 | **0.350** |
| phb003 | **0.422** | 0.000 | 0.000 | 0.000 | 0.000 | **0.282** |
| phb004 | **0.270** | 0.000 | 0.000 | 0.000 | 0.000 | **0.355** |
| phb005 | **0.186** | 0.000 | 0.000 | 0.000 | 0.000 | **0.263** |
| phb006 | **0.456** | 0.000 | 0.000 | 0.000 | 0.000 | **0.314** |
| phb007 | **0.418** | 0.000 | 0.000 | 0.000 | 0.000 | **0.388** |
| phb008 | **0.365** | 0.000 | 0.000 | 0.000 | 0.000 | **0.199** |
| scr001 | **0.494** | 0.000 | 0.000 | 0.000 | **0.163** | 0.000 |
| scr006 | **0.487** | 0.000 | 0.000 | 0.000 | **0.357** | 0.000 |
| scr007 | **0.651** | 0.000 | 0.000 | **0.210** | 0.000 | 0.000 |
| scr008 | **0.545** | 0.000 | 0.000 | 0.000 | **0.255** | 0.000 |
| sep500 | **0.462** | 0.000 | 0.000 | 0.000 | 0.000 | **0.168** |
| sep508 | **0.413** | 0.000 | 0.000 | 0.000 | 0.000 | **0.202** |
| sep509 | **0.433** | 0.000 | 0.000 | 0.000 | 0.000 | **0.226** |
| sep510 | **0.525** | 0.000 | 0.000 | **0.085** | 0.000 | 0.000 |
| sep511 | **0.310** | 0.000 | 0.000 | 0.000 | 0.000 | **0.108** |
| soc001 | **0.444** | 0.000 | 0.000 | 0.000 | 0.000 | **0.638** |
| soc002 | **0.436** | 0.000 | 0.000 | 0.000 | 0.000 | **0.557** |
| soc003 | **0.383** | 0.000 | 0.000 | 0.000 | 0.000 | **0.708** |
| soc004 | **0.449** | 0.000 | 0.000 | 0.000 | 0.000 | **0.685** |
| soc005 | **0.486** | 0.000 | 0.000 | 0.000 | 0.000 | **0.661** |
| sui001 | **0.647** | 0.000 | 0.000 | **0.185** | 0.000 | 0.000 |
| sui002 | **0.740** | 0.000 | 0.000 | **0.260** | 0.000 | 0.000 |

Figure S1 illustrates the average scores on our psychopathology dimensions as a function of the clinical groups present in our sample.

**Figure S1.**
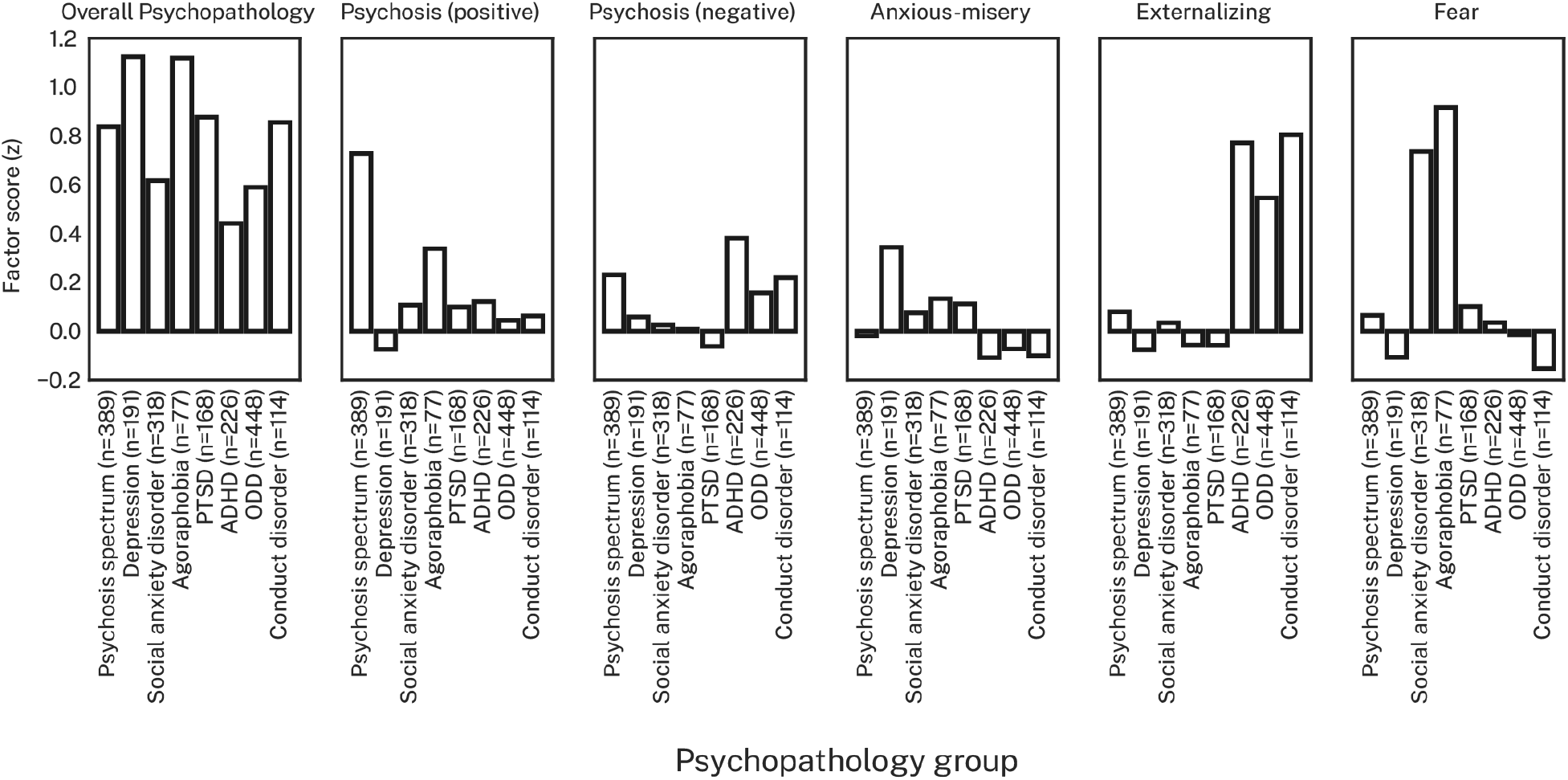
Mean psychopathology symptom dimensions as a function of psychopathology groups. Groups are the same as those presented in Table 1 in the main text. Only groups where *n*≥50 are shown here.

Figures S2 and S3 illustrate the effects of sex and age on our psychopathology dimensions. These effects were controlled for in our prediction models (see main text).

**Figure S2.**
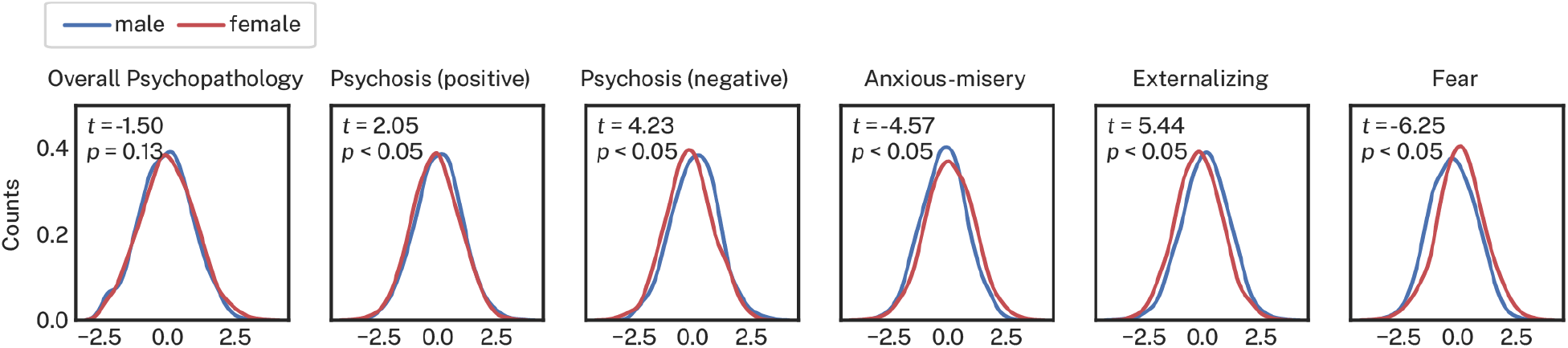
Sex effects across psychopathology dimensions in the full sample (n=1,376).

**Figure S3.**
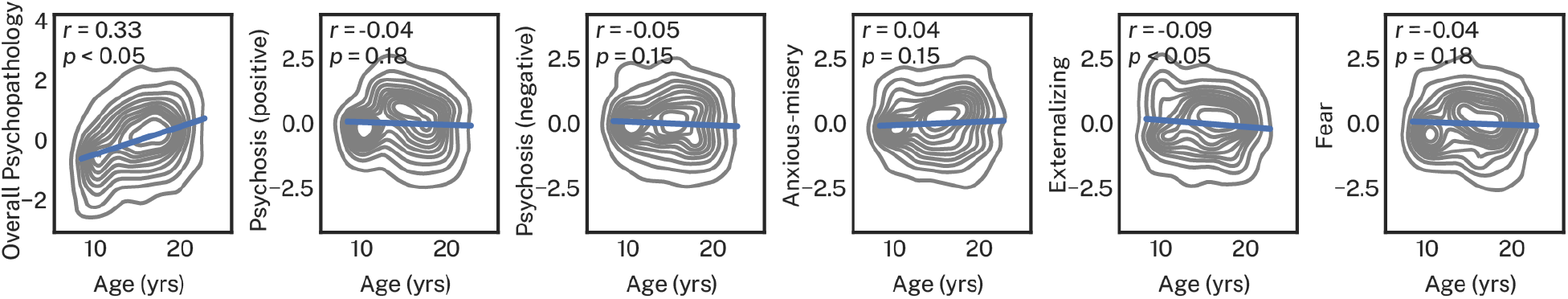
Age effects across psychopathology dimensions in the full sample (n=1,376).

### Normative models of cortical volume

For both males and females, the normative model revealed that greater age was associated with whole brain decreases in cortical volume (Figure S4A). Figure S4B shows the out-of-sample prediction error and explained variance from the normative models run for each brain region. These diagnostics demonstrate that our normative models captured the effects of age well (as a function of sex) across brain regions.

**Figure S4.**
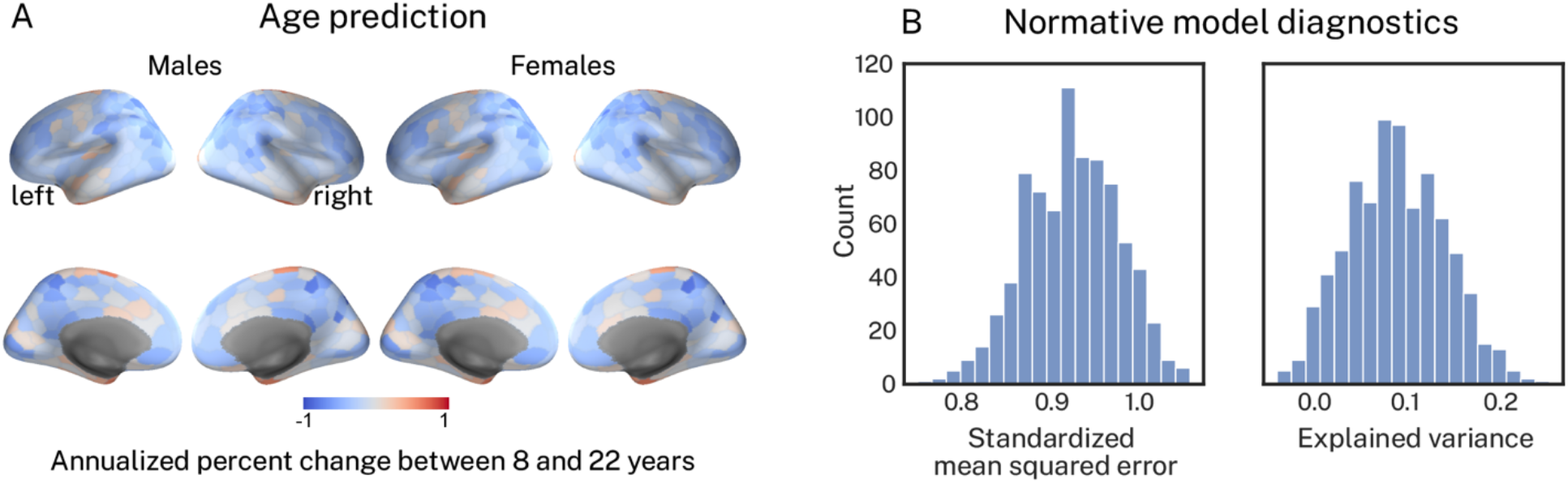
Age trajectories and model statistics from the normative model. **A**, Age trajectories for each sex learned by the normative model. **B**, Out-of-sample standardized mean squared error and explained variance from the normative models run for each region in the Schaefer400 parcellation.

### Principal component analysis of deviations from normative neurodevelopment

In the main text, prior to running our ridge regression prediction models, we reduced each of our multivariate patterns of deviations and raw cortical volume down to their respective dominant sources of variance using principal component analysis (PCA). Figure S5 shows that for both deviations and raw cortical volume estimates, variance in the data was mostly captured by the first PC. Subsequent PCs explained ~1% variance each. PCs beyond PC 9 for deviations and PC 8 for raw volume explained <1% variance each. In our regression models (see main text), we sought to retain only PCs that explained at least 1% of the variance. However, in order to maintain an equivalent number of features in our regression models, we retained 9 PCs for both deviations and raw volume. Thus, we retained 1 additional PC for raw volume that had <1% explained variance. The PC coefficients from the first 9 PCs for deviations and raw cortical volume are shown in Figure S6 and Figure S7, respectively.

**Figure S5.**
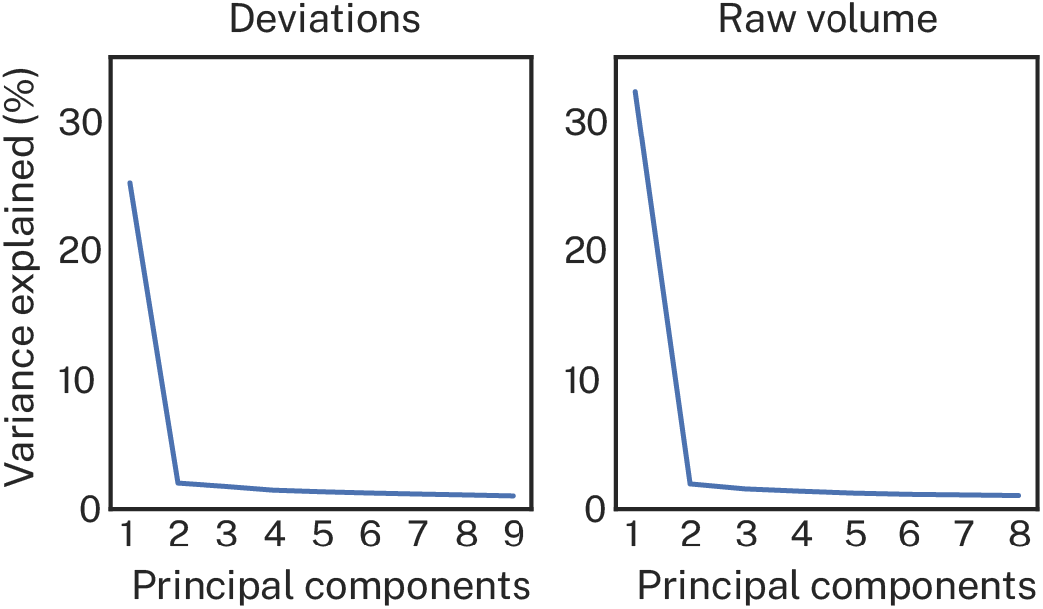
Variance explained in principal components derived from deviations and raw cortical volume.

**Figure S6.**
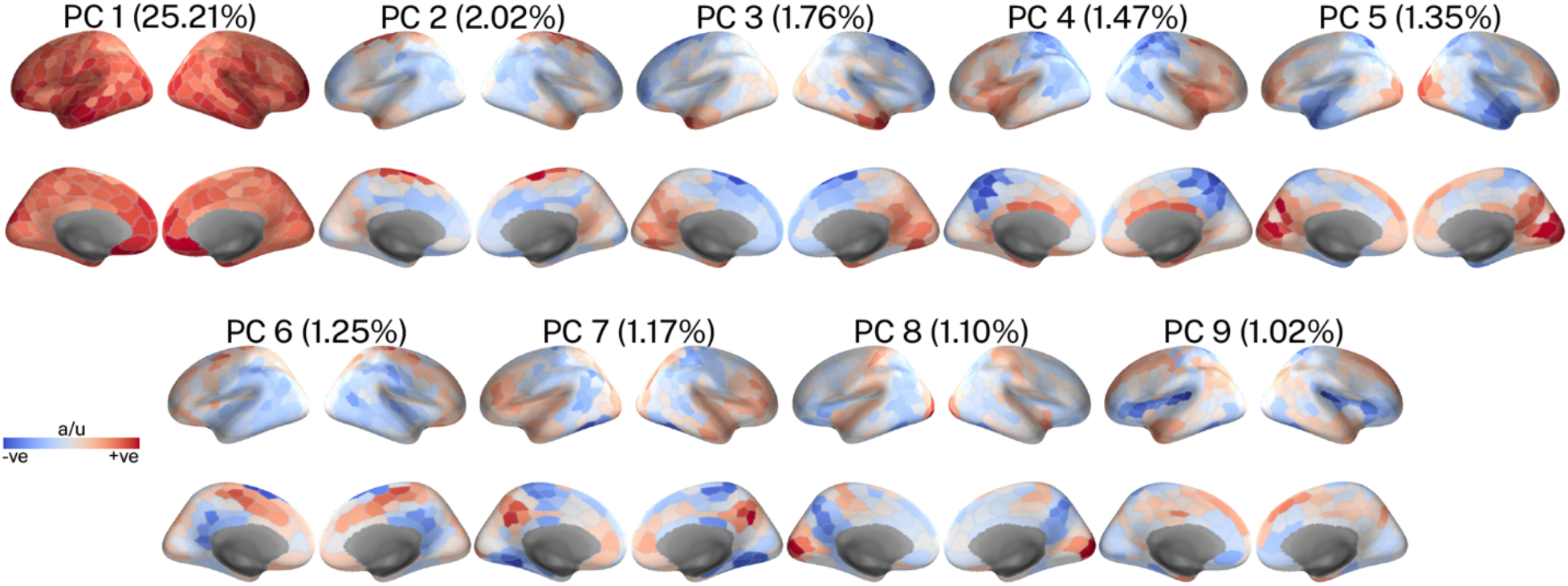
Coefficients of the principal components derived from deviations from normative neurodevelopment of cortical volume. The explained variance of each principal component is shown in parentheses.

**Figure S7.**
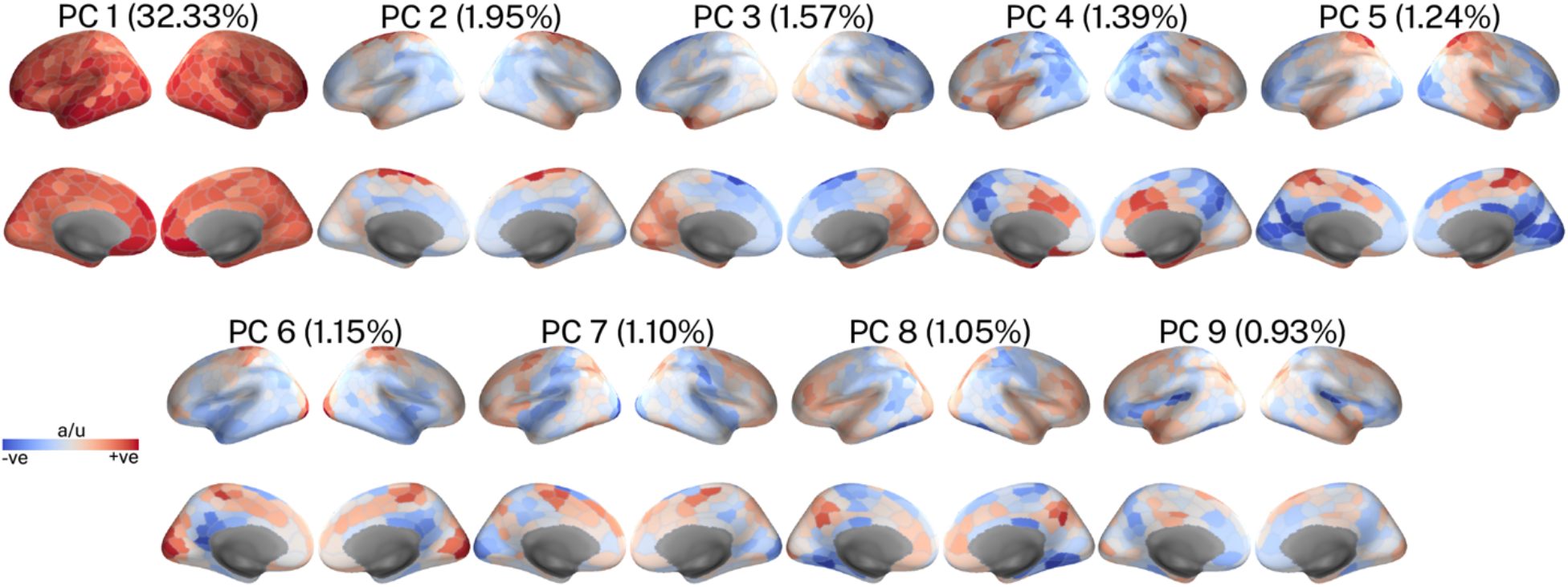
Coefficients of the principal components derived from raw cortical volume. The explained variance of each principal component is shown in parentheses.

### Psychopathology dimensions explain regional deviations from normative neurodevelopment

We compared the effect sizes of correlations between overall psychopathology and regional deviations against the effect sizes observed for specific psychopathology dimensions using bootstrapped samples. In the main text, we determined overall psychopathology to have yielded significantly larger effect sizes if the lower bound of the 99% confident interval (CI) of the bootstrapped effect size differences was greater than 0. Here, we repeated that analysis but determined the effect size differences to be significant if the lower bound of the 95% CI was greater than 0 (equivalent to *p* < 0.05, uncorrected). These results are shown in Figure S8 below.

**Figure S8.**
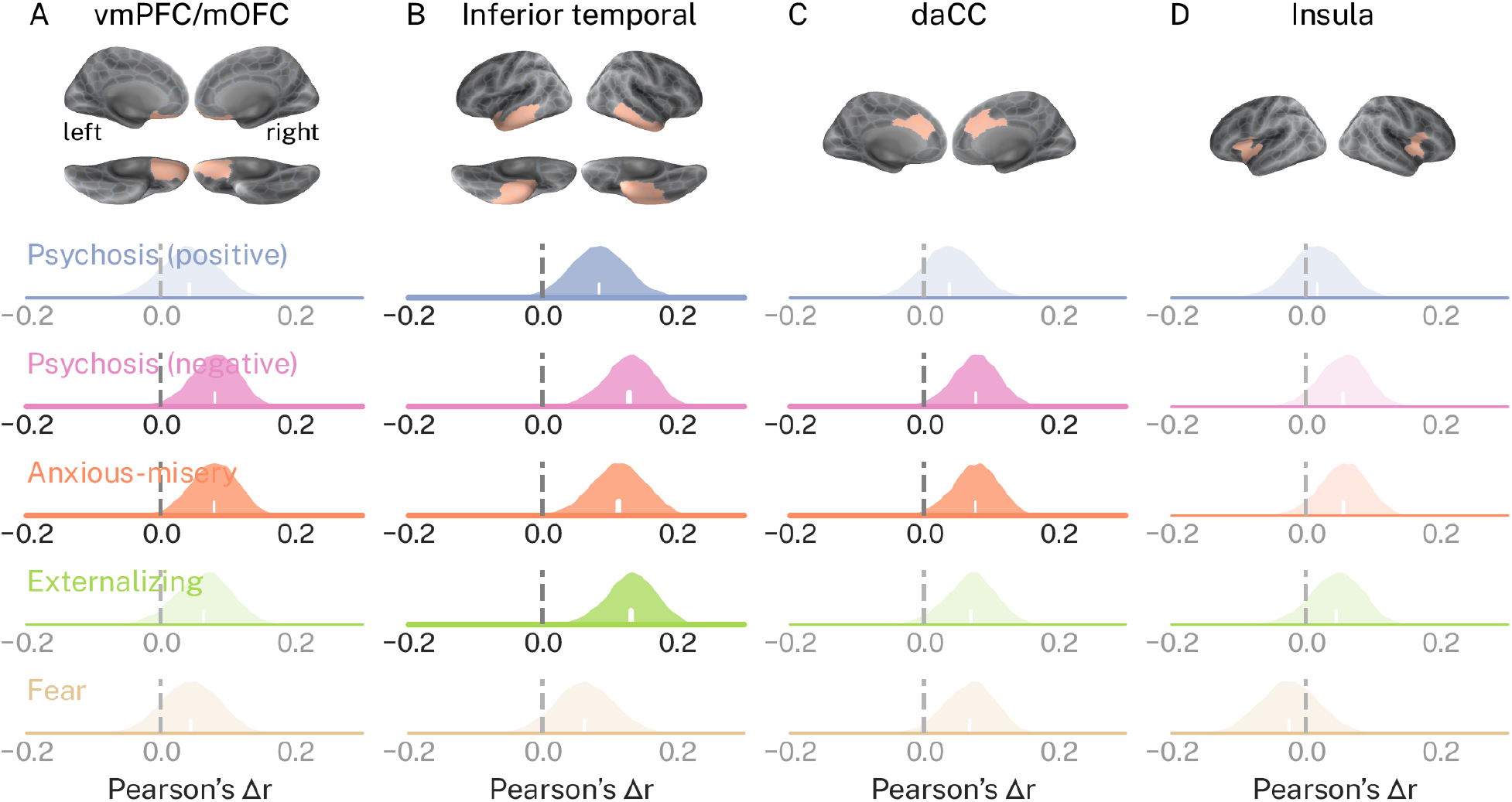
Correlations between overall psychopathology and deviations from normative neurodevelopment are stronger than correlations observed for specific dimensions of psychopathology. In each subplot, distributions of absolute Pearson’s correlations between each specific psychopathology dimension (rows) and regional deviations (columns) were subtracted from absolute correlations observed for overall psychopathology in the same region. Performing this subtraction 10,000 times across bootstrapped samples generated distributions of effect size differences, Δ*r*. Positive Δ*r* indicates that correlations for overall psychopathology were greater when compared to those observed for the specific dimensions. Δ*r* distributions for which the lower bound of the *95*% confidence interval was greater than 0 are shown with heavier stroke and no transparency.

#### Sensitivity analyses

##### Socioeconomic status

In the main text, we reported results for ridge regression prediction models that included age and sex as nuisance covariates (see Figures S2 and S3 above for sex and age effects on psychopathology dimensions). Here, we repeat these analyses, this time also including years of maternal education as a proxy for socioeconomic status (Figure S9). Figure S10 shows prediction performance largely dropped below significance when including years of maternal education as a nuisance covariate, suggesting that our proxy of socioeconomic status confounded our prediction models. Nevertheless, the general trend of deviations outperforming raw cortical volume was preserved.

**Figure S9.**
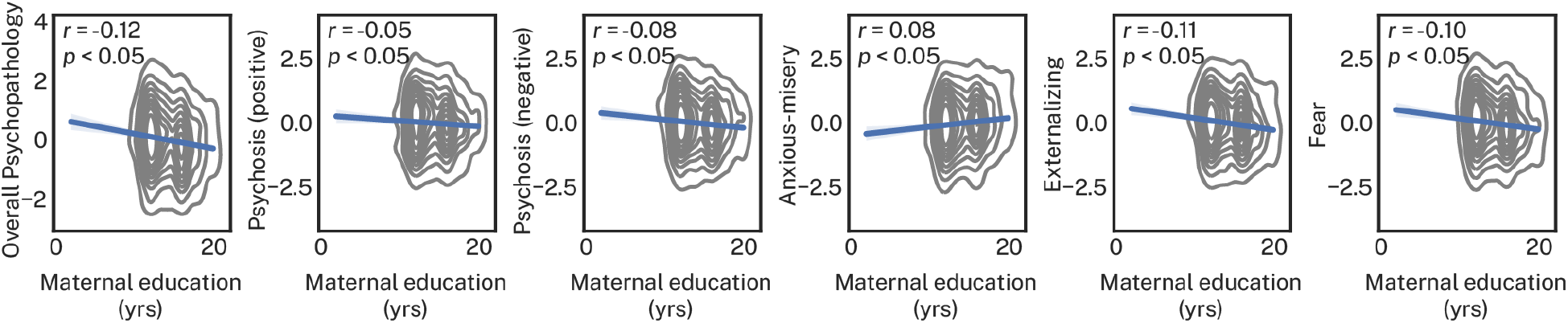
Effect of years of maternal education on psychopathology dimensions in the full sample (n=1,376).

**Figure S10.**
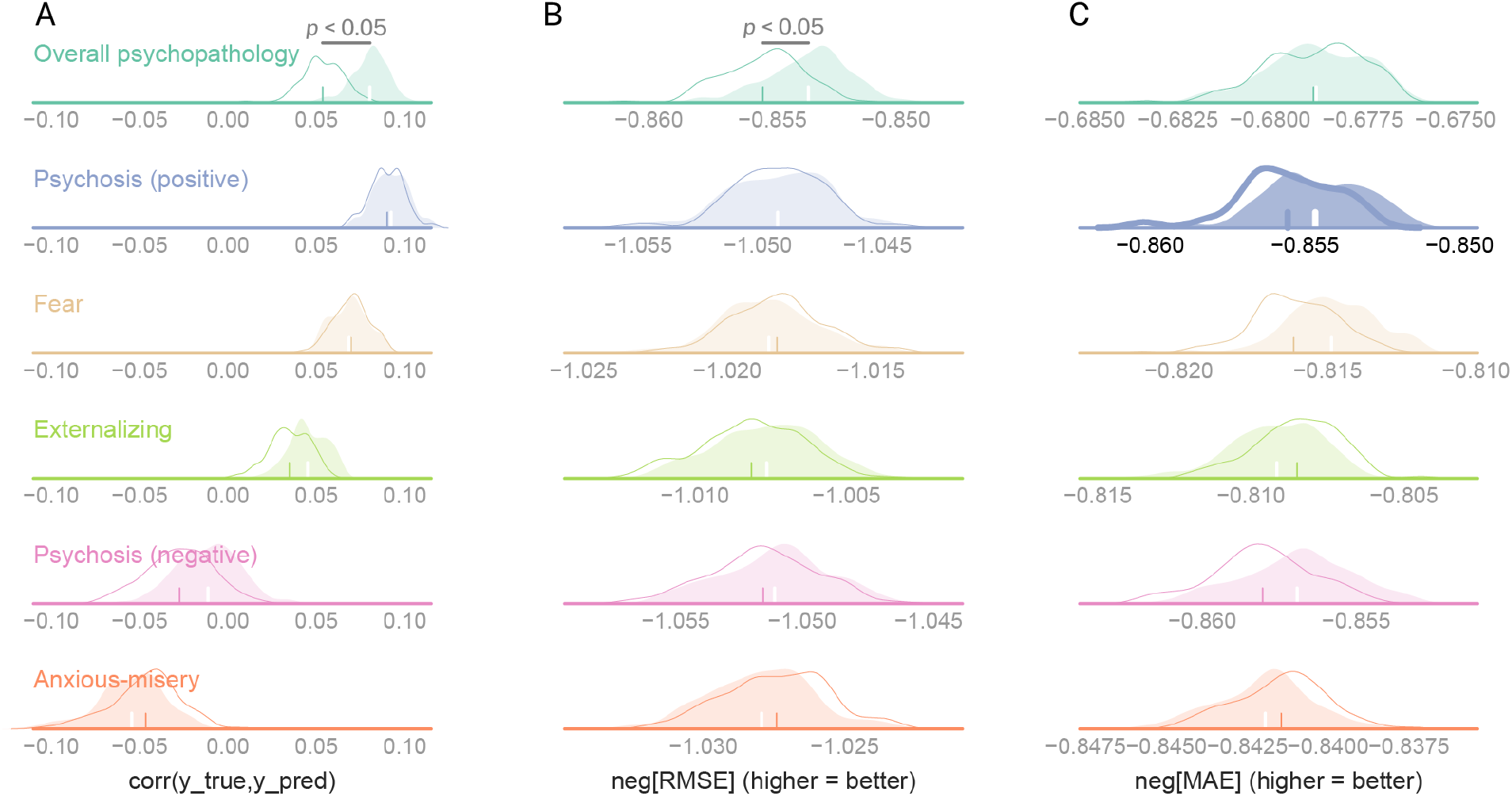
Prediction model including years of maternal education as a nuisance covariate. Predictive performance for each of six dimensions of psychopathology (rows) as a function of multiple scoring metrics (columns A-C). In each subplot two distributions are presented: one that illustrates predictive performance derived from raw cortical volume (white distribution with colored outline), and one that illustrates predictive performance derived from deviations from normative models (colored distribution). Distributions of predictive performance that did not yield above chance performance are shown with partial transparency and lighter stroke.

##### Parcellation scheme

In the main text, we reported results for the Schaefer400 parcellation. Here, we repeated our ridge regression prediction model a different parcellation known as the Lausanne atlas with 463 regions. When controlling for age and sex, we found that deviations yielded significantly higher correlations between true and predicted *y* (deviations, mean *r* = 0.087; raw volume, mean *r* = 0.064) and significantly higher negative RMSE (deviations, mean negative RMSE = −0.854; raw volume, mean negative RMSE = −0855) for overall psychopathology. Also, we found that deviations yielded significantly higher correlations between true and predicted *y* for psychosis-positive (deviations, mean *r* = 0.039; raw volume, mean *r* = 0.011) and externalizing dimensions (deviations, mean *r* = 0.054; raw volume, mean *r* = 0.039). Finally, raw volume never yielded significantly better out-of-sample predictive performance compared to deviations.

